# Femtomolar-level PCR-free quantification of microRNA cancer biomarkers in serum

**DOI:** 10.1101/2022.12.30.522268

**Authors:** Anastassia Kanavarioti

## Abstract

We developed a technology to measure microRNA (miRNA) copies in serum and tested it on a commercially available combined human serum (H6914 from Sigma-Aldrich). Copies of miR-15b and miR-16, believed to be constant among healthy and diseased individuals, were measured and agreed with the ones reported by Mitchell PS et al. (2008). Cancer biomarkers let7-b, miR-21, miR-141 and miR-375 varied 3,000 to 6,000 copies per 1 microLiter H6914 (5-10 femtomolar (fM)). Detection and quantification of oligos and miRNAs at such low concentration was shown earlier. It is accomplished by repurposing the commercial MinION nanopore platform to conduct single-molecule voltage-driven ion-channel measurements, employing osmium-tagged oligo probes, and using a publicly available algorithm. These miRNAs were quantified in the serum of healthy individuals or cancer patients using novel optimized probes and a detailed protocol that delivers miRNA copies with better than 85% confidence across all concentrations. A linear correlation, the same with healthy and cancer serum samples, is observed between miR-15b copies and RNA concentration in serum. The assay’s simplicity, readiness, sensitivity, and precision advocate for its use as a Laboratory Developed Test (LDT) for disease-screening based on miRNA dysregulation.

## Introduction

Cancer takes many forms, some very aggressive, and claims lives worldwide^1^. Early diagnosis can improve cancer outcomes by detecting malignancies early on before severe symptoms appear. Tumor biopsies are invasive and typically come at a later disease phase. Blood or urine tests, so called liquid biopsies^2^, are minimally invasive and could dramatically improve early detection. miRNAs are 18 to 25-nucleotide (nt) long RNAs and publicized as the “tiny regulators” in a seminal paper 30 years ago^3^. miRNAs were shown to exhibit regulatory function over the post-transcriptional expression of messenger RNAs, playing a critical role in physiological or non-physiological (disease) pathways^3,4^. Currently there are about 2,300 human miRNAs identified^5^. They are present in biological fluids and surprisingly stable^6,7^, rendering them useful biomarkers for cancer and other diseases^6-12^. In the last 20 years, over 140,000 peer-reviewed publications have correlated disease’s onset, progression, severity and, recurrence after treatment, with a small panel of miRNAs for many diseases, including cancers.

Copies of miRNAs may measure as low as 1,000 per 1 μl of blood (1.7 fM), which is a billion-fold less compared to concentrations required by many analytical tools. Current technologies for miRNA quantification are based on microarrays, quantitative real-time PCR, and next-generation sequencing^13-15^. These assays are complex, prone to error, require skilled personnel, expensive infrastructure, and thus unsuitable for screening and implementation at a point-of-care (POC) facility. Since miRNA dysregulation is believed to be the cause for the onset of many diseases, an easy to carry-out, reliable, and accurate miRNA assay using blood or urine could be implemented at a POC facility. Such a test may be part of a regular check-up and provide a broader picture of an individual’s health status.

In contrast to traditional analytical technologies, nanopores are well suited for trace measurements. Numerous studies document the use of experimental nanopores to detect charged chemical and biological components, including miRNAs^16-33^. In 2014 Oxford Nanopore Technologies (ONT) commercialized a device, the MinION, equipped with a consumable flow cell that carries an array of 2048 proprietary nanopores. This platform is currently used to sequence long DNA and RNA, but not short nucleic acids like the miRNAs. Sequencing is realized after library preparation, and the incorporation of a motor enzyme that guides the nucleic acid in a slow, controlled manner via a pore. Instead of sequencing, we conduct single-molecule, unassisted, voltage-driven, ion-conductance experiments. We then analyze the acquired raw ONT data, ion-current as a function of time (*i-t*), with a publicly available algorithm, (*OsBp_detect*)^34^, developed for this application. Proof-of-principle to use the MinION for quantification of oligonucleotides, including miRNAs, was reported earlier^30^.

A nanopore, solid-state or natural, embedded within an insulating membrane separating two compartments filled with an electrolyte solution, allows molecules to traverse from one side of the membrane to the other (Fig. 1a). When voltage is applied across the membrane, electrolyte ions flow via the pore(s) producing a constant ion current (*I*_*o*_). Molecules that fit through the nanopore cavity may traverse and the larger/wider ones will not. Translocation of charged molecules, including nucleic acids is expedited/driven by the voltage drop across the membrane. A traversing molecule occupies part of the space within the nanopore which reduces the available space for the electrolyte ions. Single molecule translocation reduces the constant open pore ion-current (*I*_*o*_) to a new residual level (*I*_*r*_) and creates an event/peak at *I*_*r*_ with duration (τ) equal to the time the molecule occupies the pore (Fig. 1b).

**Figure 1:**
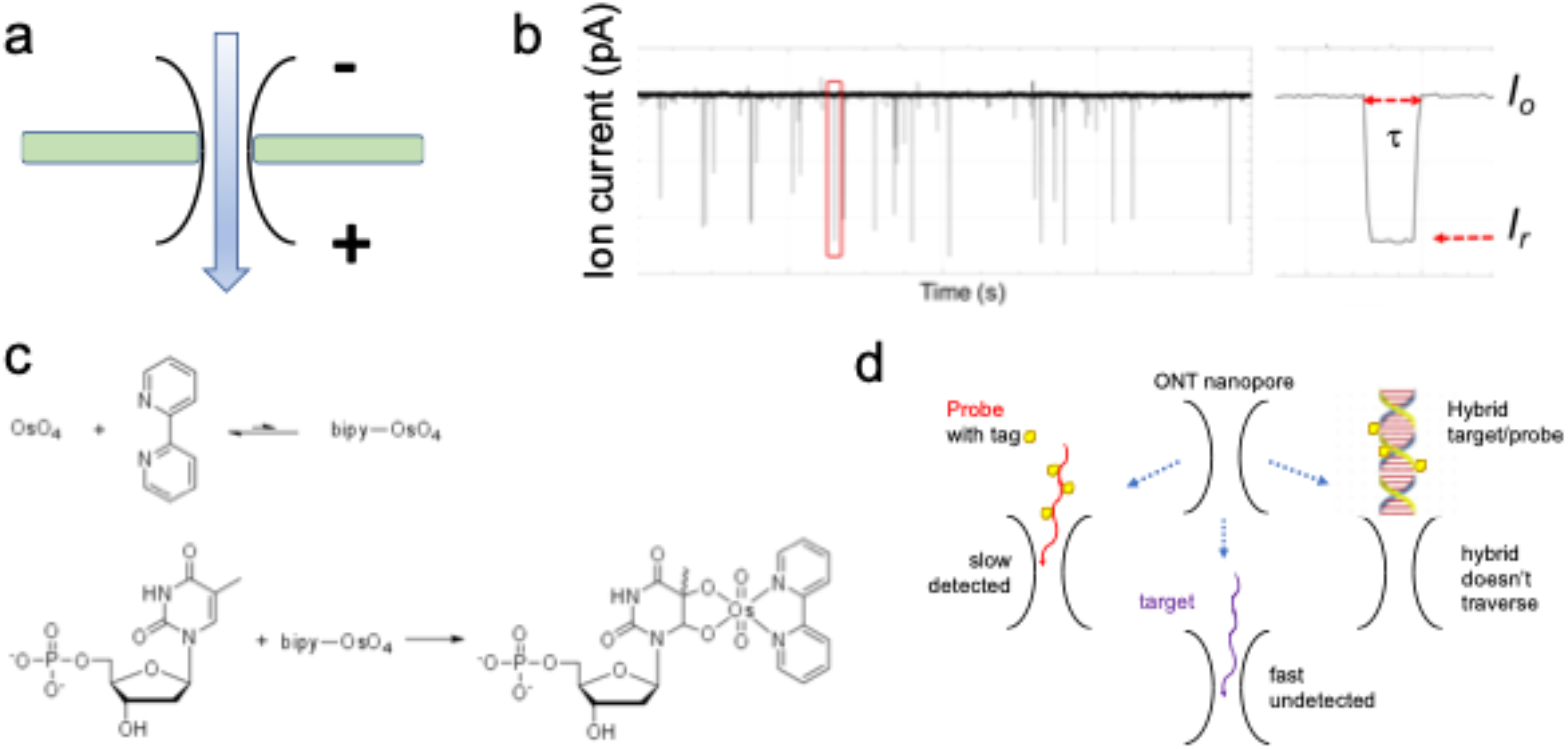
The critical parameters of the proposed diagnostic assay (with permission from reference 30). **(a)** Schematic representation of a nanopore within a planar bilayer lipid membrane that separates two electrolyte filled compartments. Applying a constant voltage guides the passage of ions through the nanopore creating a measurable ionic current (*I*_*o*_ in (b)). **(b)**The *i-t* trace obtained from a voltage-driven ion-channel experiment, with constant flow of electrolyte ions (*I*_*o*_), is interrupted by the passage of molecules that appears as events or peaks with residual ion current *I*_*r*_ and residence time τ. **(c)** The OsBp labeling reaction shown her on 5’-monophosphate-Thymidine: OsO_4_ and 2,2-bipyridine (bipy) have a low association constant, but their mixture adds to the C5-C6 double bond of pyrimidines and forms a stable conjugate, the osmylated nucleic acid. Labeling a nucleic acid using OsBp was discovered in the 1960s and has been extensively used^35-38^. OsBp adds to pyrimidines leaving the purines intact. The reactivity of OsBp towards Thymine (T) is much higher compared to Uridine (U) and Cytosine (C). Labeling was optimized to enhance T selectivity^39^ and further suppress purine inactivity^40^. Osmylation renders the oligo/probe bulkier and markedly slower compared to intact nucleic acids. The addition of OsBp creates a chromophore that absorbs around 312 nm where native nucleic acids exhibit negligible absorbance. This chromophore enabled the development of a quality control assay for use in the manufacturing of osmylated nucleic acids (see Methods). **(d)** Cartoon to illustrate the concept behind this miRNA quantification assay. Short DNAs and RNAs, including miRNAs, traverse relatively fast, a few μs per base^18^, and pass mostly undetected by the MinION platform which uses 3kHz and reports 3 data points per 1 msec. In contrast to intact oligos the osmylated oligos (probes) are detected and produce events^29,30^. When an probe is added to a sample that contains its complementary miRNA (target), the probe and the target form a hybrid that doesn’t traverse the pore. When the target’s concentration is more than the probe’s concentration, i.e., [target] > [probe], few events are observed. When [target] < [probe], many events are observed. The target’s concentration is deduced from the probe’s known concentration.

The proprietary nanopore protein of the ONT flow cell is wide enough for ss nucleic acids but not wide enough for double-stranded (ds) nucleic acids. We exploited this feature and designed a miRNA quantification assay (Fig. 1d) ^30^ as follows: for each miRNA target its complementary oligo is chosen as the probe. The probe’s sequence is extended at one end by adding a few Thymidines (T). These Ts are selectively tagged with osmium tetroxide 2,2’-bipyridine (OsBp) (Fig. 1c). OsBp is a bulky moiety that, despite its bulkiness, permits translocation of such tagged nucleic acid via several nanopores including the ONT nanopore, and its bulkiness yields preferential nanopore-based probe detection over intact nucleic acids^27-30^. In the absence of the target miRNA, the probe traverses the nanopore and produces a characteristic event/peak. In the presence of the target miRNA, probe and target hybridize, but the hybrid is too big and cannot traverse (Fig. 1d). Hybridization leads to a dramatic decrease in observed events, or “silencing”, and serves as evidence for the presence of the target. Quantification is based on the one-to-one hybridization between target and probe and the probe’s known concentration^30^.

Here, we report on a better understanding of the ONT platform, which represents the only commercially available fully integrated nanopore array, and its potential and limitations to conduct ion-conductance experiments. We propose a detailed protocol and showcase it by determining copies of miRNAs from the combined serum of healthy male individuals (Sigma Aldrich, H6914) and from serum samples of healthy individuals or cancer patients (Table S1 in the Supplementary). For the nanopore experiments we mix the RNA fraction, as obtained via an RNA isolation kit (Methods), with one or more of our probes (Table 1) and with the ONT proprietary buffer. Measurements were made in the range 3,000 to 100,000 miRNA copies per 1 μL serum (5 to 167 fM), at a better than 85% confidence across the full range. As expected, known miRNA cancer biomarkers, miR-21, miR-375 and miR-141 measure higher in the cancer samples compared to healthy samples, albeit by only 1.5 to 3.0-fold. To compensate for RNA variation, miR-15b copy number was plotted as a function of RNA (ng/μL) concentration in the serum isolate and exhibited a perfectly linear correlation including both healthy and diseased sera. Comparison of relative abundance for miR-21/miR-15b obtained from the combined serum of healthy male individuals (H6914 1^st^ lot, Table 2) and from a healthy individual H1 (H1 in Table 3, Fig. 5) yields 0.59 and 0.61, respectively. Such agreement indicates that linear relationships between miRNA copies and serum RNA may apply to other biomarkers besides miR-15b. The assay described here is simple and requires minimal infrastructure and skill. It may be carried out as a POC test, monitor the levels of a few miRNAs of an individual as part of a regular check-up, or serve as a QC assay to confirm miRNA levels determined by other means. It may serve as an LDT for miRNA validated biomarkers, represent the earliest disease-diagnostic test based on miRNA dysregulation, and materialize the promise of personalized healthcare.

**Table 1:**
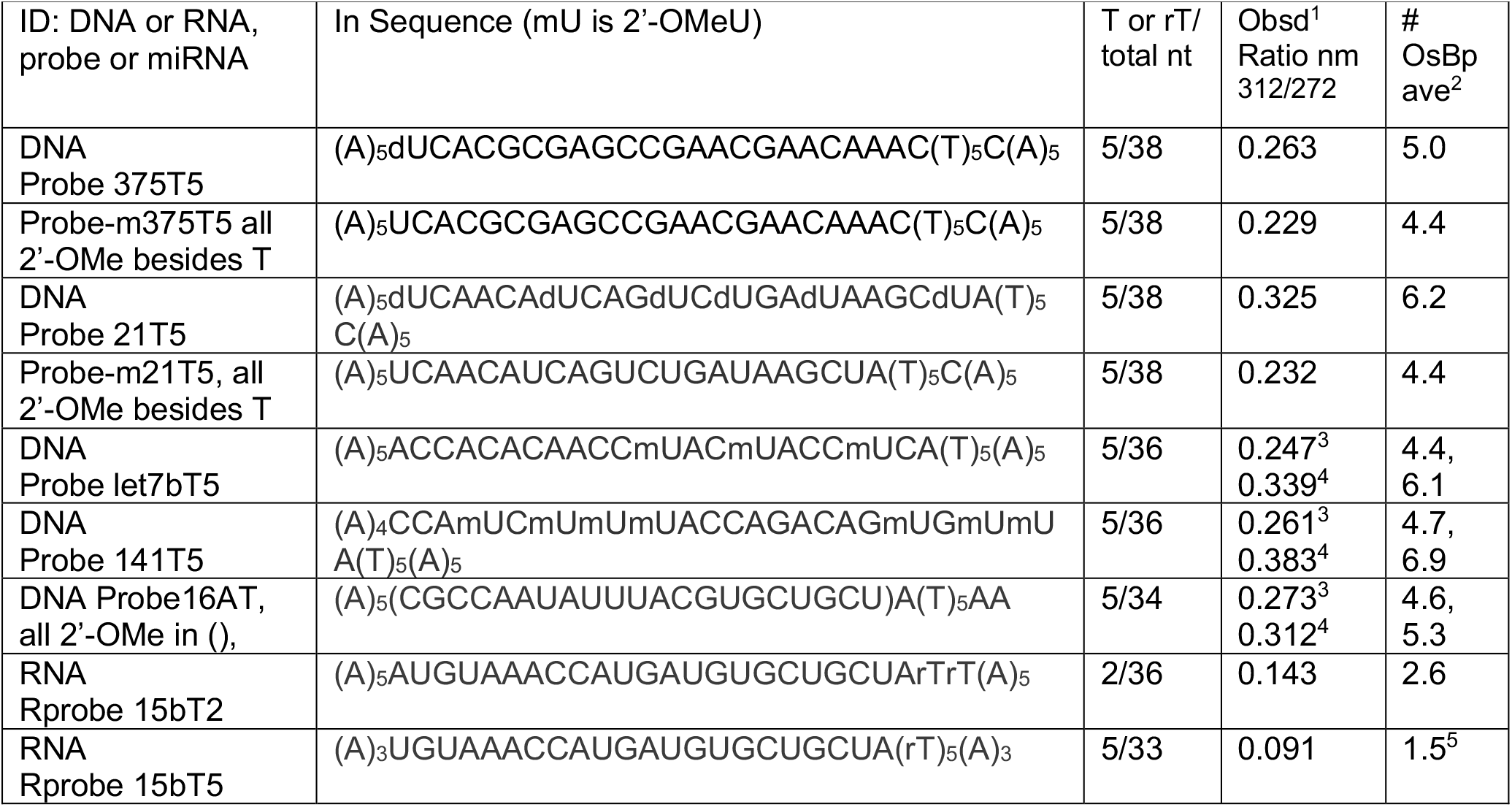
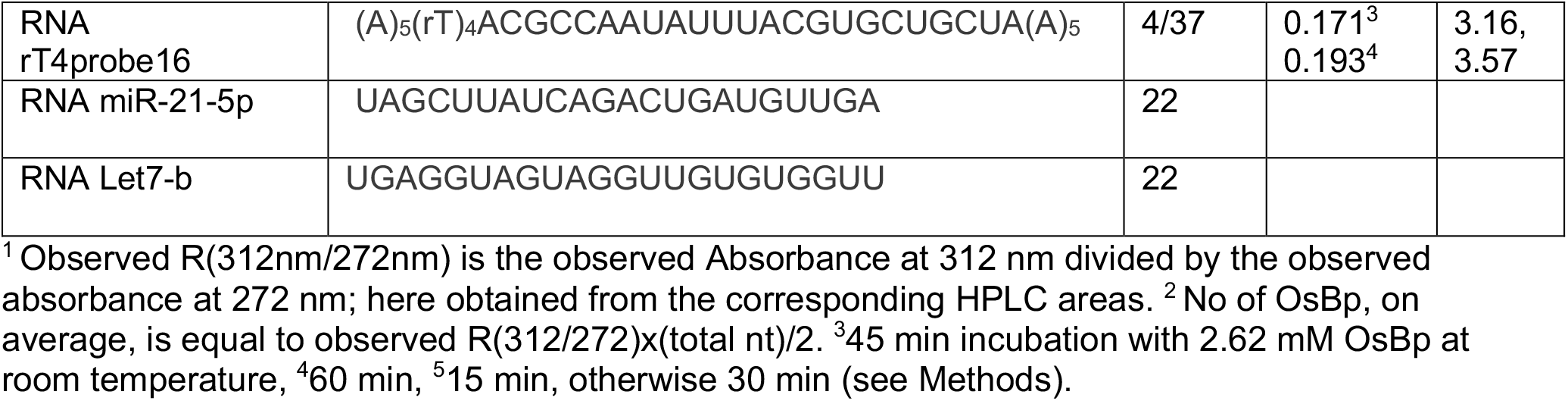
List of miRNAs, DNA or RNA oligos/probes and characterization.

**Table 2.** miRNA copies per 1 μL serum determined using the small RNA fraction.

| miRNA | H6914<sup>1</sup><br>1<sup>st</sup> lot | H6914<sup>1</sup><br>2<sup>nd</sup> lot | Pancreatic<sup>1</sup> | Breast<sup>1</sup> | Prostate<sup>1</sup> | C1<sup>1</sup> |
| --- | --- | --- | --- | --- | --- | --- |
| miR-16 | 105,125 |  |  |  |  |  |
| miR-15b | 8,855 | 8,358 | HL<sup>2</sup> | HL<sup>2</sup> | HL<sup>2</sup> | >10,800 |
| Let7-b | 6,075 |  | <2.0x HL | <1.5, 2.0x HL | 1.5 to 2.0x HL |  |
| miR-21-5p | 5,247 |  | >2.0x HL | 2.0 to 2.5x HL | 1.5 to 2.0x HL | > 2.0x HL |
| miR-375-3p | 4,620 | 4,818 | 2.0 to 2.5x HL | 1.5 to 2.0x HL | 2.0 to 2.5x HL | 1.5 to 2.0x HL |
| miR-141-3p | 3,048 |  | 1.7 to 2.3x HL | 2.3 to 2.9x HL | 2.3 to 2.9x HL | 1.5 to 2.0x HL |

<sup>1</sup> H6914 (Sigma-Aldrich, combined serum), serum samples from cancer patients (DLS, 1<sup>st</sup> batch, Table S1 in Supplementary) and from a healthy white woman (C1). Data for H6914, 2<sup>nd</sup> lot, where determined using a Monarch kit of a different lot from the one used for H6914, 1<sup>st</sup> lot. miRNA copies obtained arbitrarily as the average of two experiments, one silencing and one detection, with an apparent 7% < RSD < 16%. <sup>2</sup>HL stands for Healthy Level, as determined from H6914 1<sup>st</sup> lot.

**Table 3.** miR-15b copies per 2 μL serum (in contrast to Table 2) as a function of RNA (ng/μL) isolated from serum (see Fig. 6)

| ID<sup>1</sup> | Total RNA<br>by CE<sup>2</sup> | Copies from total RNA,<sup>3</sup><br>solid symbols |  | Copies from small RNA,<sup>3</sup><br>open symbols |  |
| --- | --- | --- | --- | --- | --- |
| | ng/μL | Healthy | Cancer | Healthy | Cancer |
| H1 | 16.2 |  |  | 16,100 |  |
| H2 | 14.4 |  |  | 13,300 |  |
| H4 | 9.1 | 9,188 |  |  |  |
| C1 | 20.7 | 21,852 |  | >21,600<sup>4</sup> |  |
| H6914, 2<sup>nd</sup> lot | 16.0 |  |  | 16,716 |  |
| H4522 | 8.4 | 7,539 |  | 7,539 |  |
| Pancreatic | 9.5 |  |  |  | 8,908 |
| Prostate CAN4 | 12.0 |  | 11,719 |  |  |
| Prostate CAN6 | 8.0 |  | 7,950 |  |  |
| Breast CAN9 | 9.1 |  | 8,922 |  |  |

<sup>1</sup> List of serum samples (DLS, 2<sup>nd</sup> batch, Table S1 in Supplementary).
<sup>2</sup> CE stands for Capillary Electrophoresis (see Methods). <sup>3</sup>Total or small RNA was obtained using the Monarch kit from NEB, using indiscriminately 1<sup>st</sup>, 2<sup>nd</sup> or subsequent lots. H1, H2, H4 are from healthy individuals, IDs for C1, Pancreatic and H6914, 2<sup>nd</sup> lot, identify the same source as in Table 2. H4522 combined serum from healthy male plasma (Sigma Aldrich). Serum samples from DLS that measured total RNA < 7 ng/ μL by CE were not tested by nanopore.
<sup>4</sup> From Table 2, adjusting for 2 μL vs. 1 μL serum.

## Results and Discussion

### Using RNA isolated from serum as the miRNA source

Experiments loading 1 or 2 μl amounts of serum mixed with the ONT buffer directly onto the MinION flow cell often led to substantial nanopore inactivation. Therefore, we used the Monarch kit from New England Biolabs (NEB No T2010S) to isolate total RNA from serum, and from the total RNA to isolate small RNA which contains RNAs less than 200 nt long including miRNAs. The kit’s protocol yields 1μL total RNA or 1μL small RNA from 2μL serum. When tested side-by-side both total and small RNA fractions gave comparable results (see later). Similarly, chemically distinct probes (see Table 1), albeit complementary to a certain miRNA, provided comparable results. Two different lots of the Monarch kit and of the H6914 serum were tested for internal assay validation. ONT flow cell versions R9 and R10 were used successfully.

### Probe design and manufacturing

Probes are manufactured from ∼38nt long RNA or DNA oligonucleotides, which were custom-synthesized and purified by the manufacturer (Table 1). The probes with the best nanopore-detecting properties are composed of a main sequence complementary to the miRNA target, extended with 5 Adenosines (A) tails at both ends. At one of the ends and between the complementary section and the A-tail, a sequence of 5 or 2 Thymidines (T) is added in the DNA or the RNA probes, respectively. Any T present within the complementary sequence of a DNA probe is replaced by a Uridine (U). Experiments to show the probe optimization efforts are not included here. Hybridization between the probe and the target miRNA in the μM range was established earlier^30^ and retested here using High Performance Liquid Chromatography (HPLC (Methods) and Fig. S1 (Supplementary)).

### Single molecule, voltage-driven, unassisted, ion-conductance experiments with the MinION

To the best of our knowledge this assay is the first to use a nanopore array for sensing^29,30^, and is unlike other analytical tools in many ways including: (a) Only a portion of the sample’s components traverse via the pore, which does not deter from target quantification (see later) (b) Due to ONT’s slow acquisition rate at 3 data points per msec many molecules that traverse the pores are not detected/reported. (a) and (b) result in preferential reporting of our proprietary probes but prohibit counting of all translocating species. (c) The ONT flow cell contains component(s) that traverse the pores and are reported (Fig. S2, Supplementary). These components cannot be subtracted out since the flow cell is being depleted of them with every new experiment and the extent of depletion cannot be estimated. (d) An increasing number of the ONT proteinic nanopores becomes inactive with use, so that the 1^st^ experiment cannot be compared to the 10^th^ experiment of the same flow cell. Issue (c) may be resolved by ONT, and (d) by manufacturers of solid nanopore arrays. Additional observations are included in the caption of Fig. S2 (Supplementary).

### Nanopore-based probe detection

This assay is best employed in comparing the results of one experiment to the results of the next experiment. For example, take an experiment conducted with ONT buffer as a sample, followed by an experiment conducted with a sample composed of a probe in ONT buffer. The second experiment will measure many more events compared to the first and showcase nanopore-based probe detection. Fig. 2a illustrates such an experiment by graphing the histogram data (Methods) acquired from running ONT buffer as a sample (blue trace), together with the data from a second experiment with a sample containing 48,300 copies of an RNA probe to target miR-15b (red trace). Another experiment (green trace), identified as “nonew”, is conducted using the already loaded sample, after letting the flow cell sit at room temperature for about 15 min. The 2^nd^ run typically exhibits a profile comparable to the 1^st^ run and splits an otherwise 1.5 h experiment into two 45 min experiments which appears to prolong flow cell life. Fig. 2a suggests that the probe at a load of 48,300 copies (red trace, 1^st^ run) exhibits 3-times more events compared to the ONT buffer and thus the probe’s presence is easily deduced. High number of events in the 2^nd^ run (green trace, (nonew)) confirms the presence of the probe which appears to take more than 45 min to translocate. Detecting 48,300 oligo copies (1.07 fM concentration in the 75μL sample/volume of the nanopore array) showcases the potential of this platform for trace analyses.

**Figure 2.**
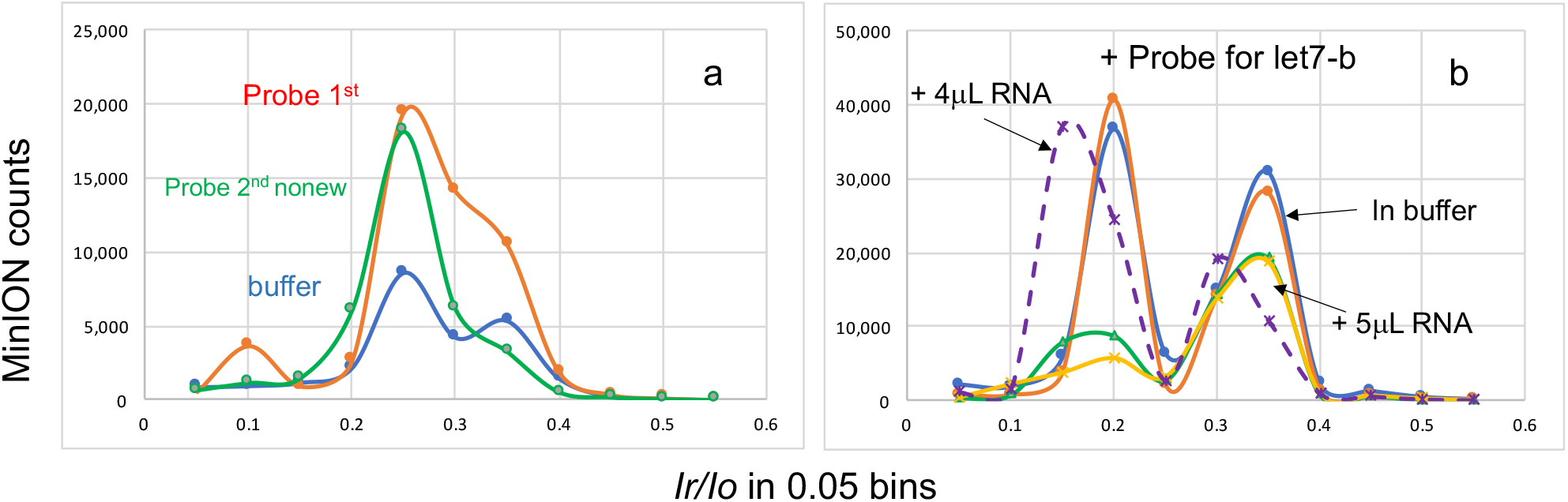
Events plots of ion-conductance experiments illustrate (a) discrimination between probe and buffer and (b) detection or silencing of probe. **(a)** Three consecutive experiments conducted with the same flow cell (FAT12944, Table S2 in Supplemental), each lasted 45min at -180mV. Data for each plot represent the histogram obtained by *OsBp_detect* analysis (Methods). ONT buffer is the control sample (blue trace) and a probe for miR-15b (Rprobe 15bT5 with 1.5 OsBp moieties, see Table 1) in ONT buffer at 48,300 copies (red trace). An additional run using the same sample was conducted at the same conditions (2^nd^ run, green trace (nonew)). Compared to the events from the ONT buffer sample, both runs of the probe in buffer exhibit about 3-times more events at (*I*_*r*_*/I*_*o*_)_max_ ≅ 0.25, which is a probe feature. The observation that not all the probe molecules traverse within the first 45 min is not uncommon. **(b)** Three samples were tested in consecutive 45 min experiments conducted at -180mV using the same flow cell (FAT43745, Table S2 in Supplemental). Each sample contained 54,000 copies of probe let7bT5 (Table 1) to target let7-b in H6914, 1^st^ lot. The first sample contained the probe in buffer, a 1^st^ run was conducted (blue trace), followed by a 2^nd^ run with no sample addition (red trace (nonew)). The second sample contained the probe and 5μL small RNA from H6914 (green trace), followed by a 2^nd^ run (yellow trace (nonew)), and exhibited dramatically fewer counts (silencing) compared to the first sample with the probe only. The third sample contained the probe and 4 μL small RNA from H6914 (purple dashed trace) and exhibited dramatically more counts (detection) compared to the sample with 5 μL small RNA. The difference between 4 and 5uL small RNA cannot explain the 4-fold increase in counts, as documented in Fig S3a (Supplementary). One concludes 10,800 < let7-b < 13,500 copies per 1 μL of small RNA from serum H6914, from 54,000/5 and 54,000/4, respectively, or 5,400 < let7-b < 6,750 copies per 1 μL of serum H6914 due to the 2-fold concentrating effect of the RNA purification step (see Methods). The fact that we estimate miRNA copy number with better than 85% confidence, and conjecture this from experiments that differ often by over 200% in counts deserves an explanation. Presumably a 10 to 15% excess of probe or hybrid is what tilts the balance from detection to silencing, as seen in Fig. 2b and in the additional examples presented here. It is understandable that this 10 to 15% excess of probe or hybrid shifts the material from the early(*I*_*r*_*/I*_*o*_)_max_ to the late(*I*_*r*_*/I*_*o*_)_max_. The observation that this small percent of material shift leads to over 200% change in actual events is attributed to an additional large effect coming from the presence of the hybrid that “shields” the nanopores (see Discussion in text).

### Probe detection or silencing in H6914 serum

A second type of experimentation compares consecutive experiments using a probe at a constant number of copies in a mixture with an RNA aliquot at two different concentrations. Figure 2b illustrates such a comparison. In this set of experiments the probe for let7-b at 54,000 copies was tested in ONT buffer (traces, blue and red (nonew)). The same load of probe in the presence of 5 μL small RNA fraction from H6914 1^st^ lot (green, and yellow (nonew)) strongly suggest silencing due to the markedly reduced events (principle described in the caption of Fig. 1d). Conversely, in the presence of 4 μL small RNA (purple dashed trace) detection is concluded due to number of events comparable to the earlier experiment with probe only. Evidently, 4μL small RNA fraction does not contain enough let7-b copies to fully hybridize with the probe (let7-b < 54,000/4=13.500), whereas 5μL RNA, at 25% more content, contains sufficient copies of let7-b (let7-b > 54,000/5= 10,800). It is unclear what ratio of hybrid/free probe yields a silencing or a detection experiment. We arbitrarily take the average from the silencing and from the detection experiment and post it as copies let7-b = 12,150. Considering that RNA purification yields a 2-fold concentration (see Methods), then let7-b measures 6,075 copies in 1μL H6914 serum. Additional experiments with H6914 are illustrated in Figures S3 and S4 (Supplementary).

As seen in Fig. 2b and later figures the events profile exhibits two (*I*_*r*_*/I*_*o*_)_max_, one in the range of 0.3< *I*_*r*_*/I*_*o*_ <0.4 (late) attributed primarily to the ONT component(s), and the other in the range of 0.15< *I*_*r*_*/I*_*o*_ <0.25 (early) attributed primarily to our probes and the serum RNA. The observation of fewer events, typically at more than 50%, in a “hybridization or silencing” experiment is attributed to the absence of free probes to traverse, but also to the presence of the hybrid, which is attracted to the pores by the applied voltage, doesn’t traverse, but lingers around the probes, “shields” them, and prevents a large portion of all detectable species from traversing the pores. Besides the fewer number of events observed in a silencing experiment compared to a probe only experiment, the events at the early (*I*_*r*_*/I*_*o*_)_max_ decreases more compared to the events at the late (*I*_*r*_*/I*_*o*_)_max_, and thus a higher value of the Ratio = late(*I*_*r*_*/I*_*o*_)_max_ /early(*I*_*r*_*/I*_*o*_)_max_ suggests silencing, even when the total number of events remains comparable. Conversely, a lower value of this Ratio indicates detection. Change in the observed Ratio results from the preferential detection of the OsBp-tagged probes (early (*I*_*r*_*/I*_*o*_)_max_), when these probes are present and their copy number is not negligible compared to the total number of detectable species, let us say at or above 25%.

### Quantification of miRNAs from the serum of cancer patients

Figure 3 presents a consecutive series of five experiments (3 samples) conducted using small RNA isolated from the serum of a patient with pancreatic cancer (Tables 2, 3, S1 and S2 in Supplementary). The first sample contains a probe for miR-375 and sufficient small RNA load to target 9,173 miR-375 copies (blue trace, (nonew)). Very few events are observed, and silencing is concluded with miR-375 > 9,173. The conclusion of silencing is based on the comparison with an earlier sample, not shown here. The second experiment is conducted with the exact same load of probe for miR375 and sufficient small RNA load to target 11,634 copies of miR-375. Both the 1^st^ run (red trace) as well as the 2^nd^ run (green trace, (nonew)) clearly suggest detection of the probe with miR-375 < 11,634 copies per 1μL of serum (see figure caption). We chose the small RNA loads of these two experiments, so that the target miR-375 is equivalent to 2.0-fold and 2.5-fold the miR-375 copy number determined from H6914 1^st^ lot. Thus miR-375 in the pancreatic cancer sample is 2.0-to 2.5-fold overexpressed compared to the healthy level (HL) found in H6914. Fig. 3b uses as control/baseline the last run (nonew) to target 11,634 miR-375 copies (blue trace in Fig. 3b, was green trace in Fig. 3a). Fig. 3b presents an experiment (1^st^ run, red trace and 2^nd^ run, green trace (nonew)) conducted with probe for miR-141 and sufficient small RNA from the pancreatic cancer sample to target 5,289 miR-141 copies. Dramatically fewer events are observed indicating silencing in both runs and yielding miR-141 > 5,289 copies in 1μL serum. Sample to sample carry-over is estimated in the order of 10 to 15%, and this is not an issue here as the effects are in the order of 2-fold or higher. Additional nanopore experiments using RNA from the serum of cancer patients with breast or prostate cancer are shown in Fig. S5 (Supplementary).

**Figure 3:**
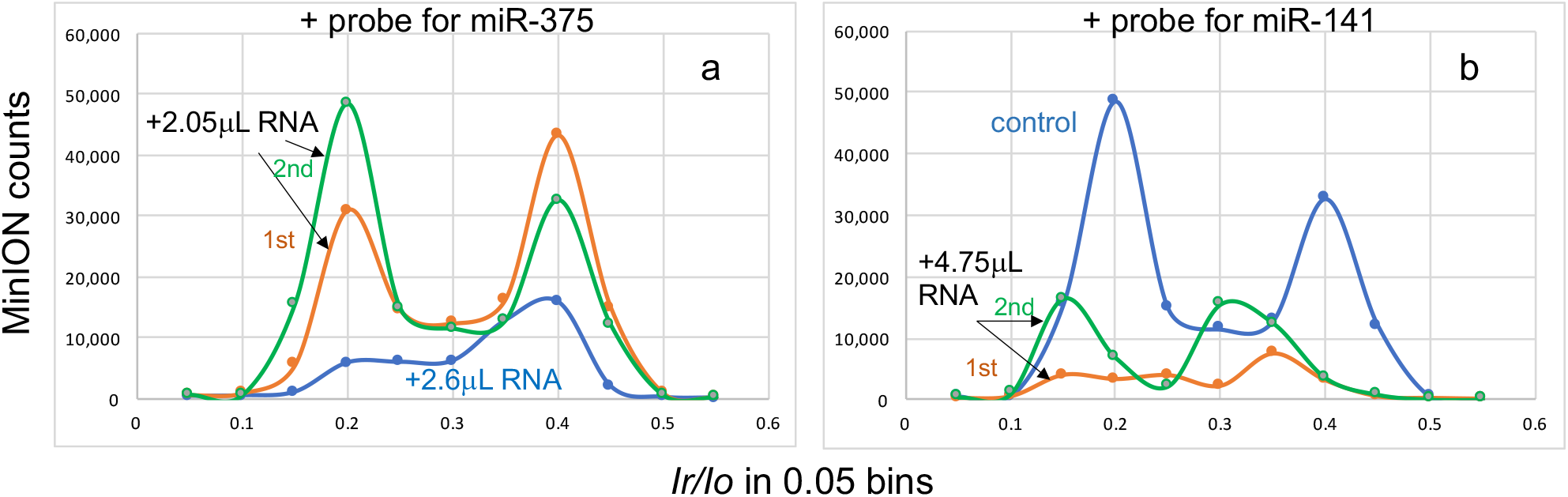
Events plots, experiments targeting (a) miR-375 and (b) miR-141 in small RNA isolated from the serum of a patient with pancreatic cancer (Tables 2, 3, S1 (Supplementary). These two figures represent a consecutive series of five 45-min long experiments conducted with flow cell FAT94165 at -180mV. The control sample (blue trace) in Fig. 3a is a “nonew” run with a sample containing 47,700 copies of probe m375T5 and 2.6 μL of pancreatic small RNA. This experiment was identified as silencing experiment (Table S2) with miR-375 > 9,173 from 47,700/2×2.6=9,173 in 1 μL serum. The next experiment was done with the same load of probe but with less RNA, i.e., 2.05 μL from the pancreatic small RNA (red trace) and exhibits 5-fold and 3-fold more events at (*I*_*r*_*/I*_*o*_)_max_ =0.2 and (*I*_*r*_*/I*_*o*_)_max_ =0.4, respectively. These dramatic increases identify a detecting experiment with miR-375 < 11,634 copies from 47,700/2×2.06=11,634 per μL serum. The following run (green trace, nonew) exhibits a similarly high number of events and confirms free probe detection. In Fig. 3b, the blue trace is the experiment presented in Fig. 3a as green trace, i.e., a detection experiment. The next (4^th^) experiment in this series is done with 4.75 μL of pancreatic small RNA and probe 141T5 at a load of 50,250 copies to target miR-141 (red trace). Total counts in this 4^th^ experiment are very low, about 1/6 of the events reported for the control, and strongly suggest silencing, which is further confirmed by the comparably low events of the corresponding 2^nd^ run (green trace, (nonew)). The conclusion is that PA miR-141 > 5,289 copies per 1 μL pancreatic cancer serum from 50,250/2×4.75=5,289.

A couple of observations are worth noting in Figures 3a and 3b. These experiments, three different samples, are conducted with the same flow cell and around the middle of its life span; the number of working channels starts at 75% and measures 54% with the last run. The overall decrease in nanopores is gradual and cannot account for the approximate 10-fold increase observed at (*I*_*r*_*/I*_*o*_)_max_ =0.2 between 1^st^ and 2^nd^ sample and the approximate 10-fold decrease observed at (*I*_*r*_*/I*_*o*_)_max_ =0.2 between 2^nd^ and 3^rd^ sample.

Another observation relates to the changes observed at the late (*I*_*r*_*/I*_*o*_)_max_ =0.4. As mentioned earlier the peak at (*I*_*r*_*/I*_*o*_)_max_ =0.2 is mostly RNA and probe, whereas the peak at (*I*_*r*_*/I*_*o*_)_max_ =0.4 is primarily due to the ONT component(s). The latter yields a large peak, as the one observed in the red and green traces of Fig. 3a (the blue trace in Fig. 3b is the green trace in Fig. 3a). In Fig. 3b the peaks shift from 0.4 to 0.3 and from 0.2 to 0.15, respectively; such shifts are not uncommon. As expected, silencing dramatically reduces the events under the early (*I*_*r*_*/I*_*o*_)_max_ peak. Surprisingly, silencing also decreases the late (*I*_*r*_*/I*_*o*_)_max_ peak. As discussed earlier this decrease is attributed to the presence of the hybrid which is negatively charged, driven towards the nanopores, doesn’t traverse due to its size, but its presence in the proximity of the nanopores prevents other molecules, including the ONT component(s), from traversing and being detected at the late (*I*_*r*_*/I*_*o*_)_max_. On one hand, the “shielding” of the nanopores by the hybrid prevents “sample titration” and direct quantitation, and on the other hand, yields a remarkable 5 to 10-fold decrease in the number of observed events between samples that differ no more than 20% in RNA load (small RNA at 2.6 and 2.05 μL) and less than 5% in probe load (47,700 vs. 50,250 copies, see caption in Fig. 3). While this assay provides a YES/NO answer regarding the presence of the target miRNA, the experimental design determines the precision by which the target is quantified. Both RNA and probe load in a sample may vary in the range of 1 to 10μL and differ by 0.1μL which leads to a dramatically small (1%) difference between two experiments regardless of the targeted copy number. We aimed at experiments with a 15% difference in one of the two components. For example, testing a constant probe count with a 7μL or an 8μL RNA sample, whereas one experiment exhibits detection and the other silencing, yields a higher than ±85% confidence, substantially higher compared to currently used technologies^13^.

### Using the total RNA fraction from serum

There are many advantages in using serum directly and it is plausible that a serum/blood filtering process will be developed later to facilitate direct testing of miRNAs in serum or blood. Similarly, it is advantageous to test total RNA isolated from serum compared to small RNA because of a shorter purification process and the fact that many RNA purification kits do not yield a small RNA fraction. We used the Monarch kit from NEB which provides access to both the total RNA and the small RNA fraction. Initially the small RNA fraction seemed like a better approach since total RNA was presumed to overwhelm the observed events. Most of the experiments were conducted with the small RNA fraction. Development of the improved probes (Table 1) enabled testing of both the small and the total RNA fractions and illustrated excellent agreement which speaks volumes for this specific purification kit and validates total RNA as test article.

Figure 4 illustrates experiments using total RNA isolated from the serum sample of a second patient with prostate cancer (ID, prostate CAN6 in Table 3). Experiments were conducted as 45-min long at -180mV. An experiment with a sample was conducted (1^st^ run), followed by a second experiment (2^nd^ run, nonew) with no sample addition. For simplicity the histogram data from the 1^st^ and the 2^nd^ runs were combined and graphed together representing a 1.5 h experiment. The first sample contains only 5.6μL total RNA, the second sample contains the same amount of total RNA and 42,000 counts of the probe for miR-15b. The third sample contains the same load of probe but less total RNA at 5.0 μL. The blue trace always represents the “control” experiment. It is easy to conclude that Fig. 4a illustrates a silencing experiment as the addition of the probe to RNA yields fewer events instead of more events. On the contrary Fig. 4b illustrates a detection experiment; less RNA (going from 5.6 to 5.0 μL) includes less miRNA target and yields more events because the probe load remains the same. One determines 7,500 < miR-15b <8,400 (see Fig. 4 caption) and miR-15b = 7,950 per μL of total RNA. Additional experiments discussed later validate the use of total RNA.

**Figure 4:**
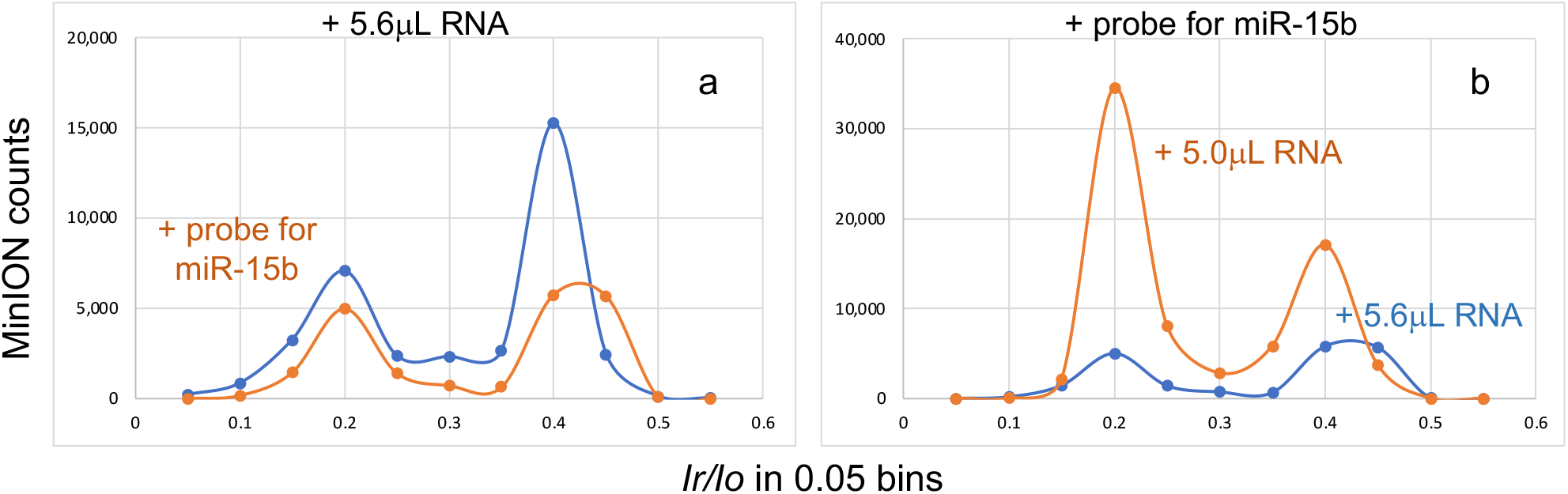
Events plots from MinION experiments targeting miR-15b in total RNA isolated from the serum of a patient with prostate cancer. See under prostate-CAN6 in Tables 2 and S1 (Supplementary). Experiments were conducted for 45min at -180mV, 1^st^ run after adding the sample, 2^nd^ run (nonew) (FAU69853 in S2 (Supplementary)). A duplicate experiment using FAU70015 yielded another set of data, but the same result (Table S2, Supplementary). For simplicity the histogram data from *OsBp_detect* analysis of both runs were combined and presented here as one 1.5 h long experiment. **Figure 4a:** 5.6μL total RNA (prostate CAN6) as control sample (blue), followed by an experiment with a mixture of 5.6μL total RNA (PRO-CAN6) and 4μL probe15bT5 at 42,000 counts (red). Despite the presence of the probe, there is only about half of the events compared to the control, consistent with silencing at this level of RNA, yielding miR-15b > 7,500 from 42,000/5.6 in the total RNA aliquot. **Figure 4b:** In this experiment the control (blue) is the earlier experiment (red trace in Fig. 4a) and it is compared to an experiment with a lower amount of total RNA at 5μL while keeping the probe counts the same at 42,000. The observation of 350% more events with less RNA clearly suggest probe detection with miR-15b < 8,400 from 42,000/5. Arbitrarily we take the average of these two relationships as miR-15b = 7,950 per μL total RNA, or miR-15b = 3,975 per μL serum.

### Using version R10 (newer) ONT flow cell

Currently ONT commercializes two flow cell versions, R9 with one constriction zone and R10 with two. Both were tested and work well with this application. Most of the earlier experiments in this study were conducted using flow cell version R9 and the small RNA fraction (see Methods). The newer version (R10.4.1 or R10) exhibits substantially more events compared to the older R9 version, due to the ONT component(s), and requires more washing with ONT buffer to deplete these components. Attempts were made to use flow cells after multiple “washing” with ONT buffer and considerable depletion of the ONT component(s), but these flow cells did not perform well. Using the R10 flow cell we demonstrate that, if the probe copies used in an experiment are close to or above 25% compared to the detected events from an ONT buffer test (see blue trace in Fig. 2a), then reliable results are obtained. Figures 5a and 5b showcase experiments to quantify miR-21 from the small RNA fraction of a healthy individual (H1) using the R10 flow cell.

**Figure 5:**
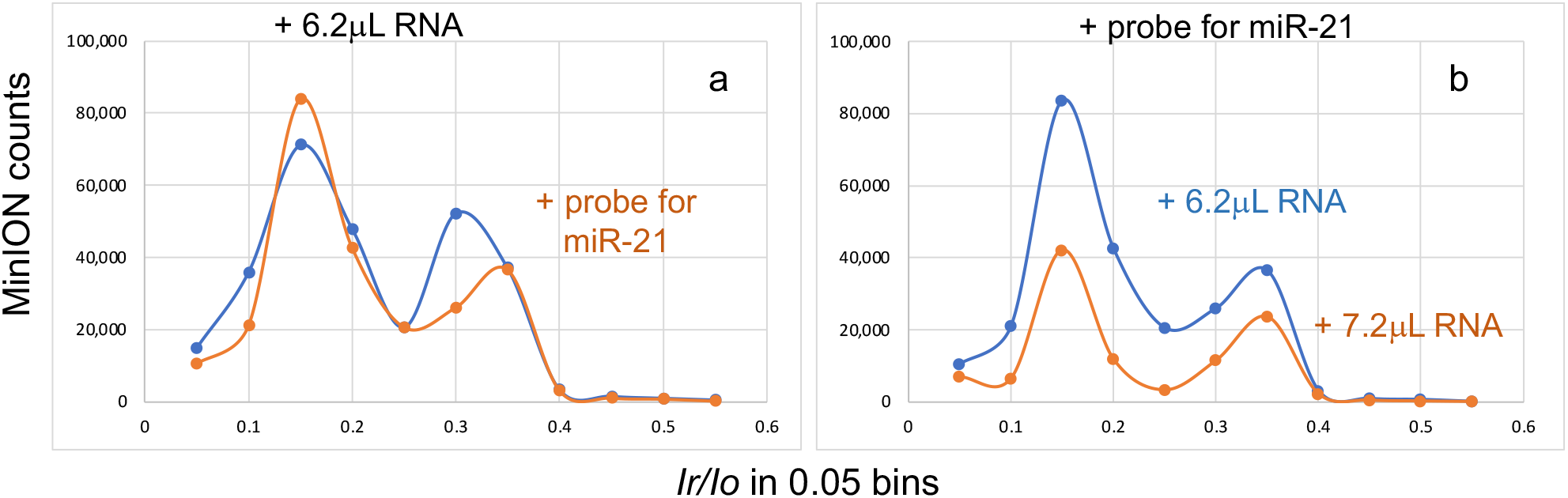
Events plots using version R10 flow cell and targeting miR-21 in small RNA isolated from the serum of a healthy individual (H1) (H1, Table S1 (Supplementary)). Experiments were conducted for 45min at -180mV with flow cell FAV45860, 1^st^ run after adding the sample, 2^nd^ run (nonew) without opening the flow cell. For simplicity the histogram data from *OsBp_detect* analysis of both runs were combined and presented here as one 1.5h long experiment. **(a)** The control sample is small RNA from the serum of a healthy individual (H1) at 6.2μL (blue trace) followed by a sample that contains the same amount of RNA from H1 with the addition of 4μL probe m21T5 at 65,040 counts (red trace). The number of events detected from these two experiments are comparable, but the Ratio late/early (*I*_*r*_*/I*_*o*_)_max_ decreases which indicates probe detection, as discussed in the text, yielding miR-21 < 10,490 from 65,040/6.2. **(b)** Here the control (blue trace) is the earlier experiment (red trace in Fig. 5a) and it is compared to an experiment with an increased amount of total RNA at 7.2μL but keeping the probe counts the same at 4μL. Dramatically fewer events as well as an increase in the Ratio late/early (*I*_*r*_*/I*_*o*_)_max_ suggest silencing at these conditions with miR-21 > 9,034 from 65,040/7.2. Arbitrarily we take the average of these two relationships as copy number miR-21 = 9,762 per μL from small RNA, or miR-21 = 4,881 per 1 μL of serum.

Just as described for the experiments in Figure 4 (flow cell vs. R9), the experiments shown in Figure 5 (flow cell vs. R10) were also conducted as two 45-min long at -180mV; the histogram data are added for simplicity and graphed together. Three samples were tested, the first used as control and contains only RNA, i.e., the small RNA fraction at 6.2μL obtained from the serum of a healthy individual (H1). The second sample contains the exact same amount of RNA and 65,040 probe copies to target miR-21. The third sample contains the same probe load but more RNA (7.2 μL). In Fig. 5a the number of events is comparable between these two samples, but probe detection is deduced from the shift of the material to the early (*I*_*r*_*/I*_*o*_)_max_. On the contrary, Fig. 5b yields a silencing experiment because the sample in the presence of more RNA (7.2 μL) exhibits about half of the counts compared to the sample with less RNA (6.2 μL). The conclusion is reached that 9,034 < miR-21 < 10,490 or miR-21 = 9,762 per μL small or total RNA in H1. Additional experiments with H1 serum to quantify miR-15b yielded miR-15=16,100 (see Table S2 under flow cell FAS88208 in Supplementary). Comparison of relative abundance for miR-21/miR-15b obtained from H6914 (1^st^ lot, Table 2) and from H1 yields 0.59 and 0.61, respectively. This remarkable agreement among experiments showcases a platform for highly accurate miRNA quantification.

### miRNA biomarkers in cancer samples (Table 2) measure higher compared to the values obtained from H6914

Table 2 compiles measurements of miRNA copies from the combined serum of H6914, and selected tests from a second/newer lot of H6914 using a different lot of the Monarch kit. Reproducibility was confirmed based on the observed comparable copy numbers for miR-15b and miR-375. Table 2 also lists the results of experiments conducted using serum from three individuals, each diagnosed with prostate, pancreatic and breast cancer at an early stage and from a healthy individual (C1). These experiments were designed to assess the expression level of cancer biomarkers compared to the level observed with H6914, and to determine how many folds higher are these biomarkers in cancer samples compared to the healthy samples. It was found that while let-7b level is less than 1.5-fold compared to H6914, the other four miRNAs tend to measure 2 or 3-fold compared to H6914. Not surprisingly the levels of these four biomarkers in C1 also appear to be less than 2-fold compared to H6914. These data confirm the notion that miRNA cancer biomarkers, like the ones tested here, are elevated in cancer. Specifically, the data agree with the notion of overexpressed levels for miR-21, a highly agreed upon biomarker for both prostate^41^ and breast^42^ cancers, as well as with the notion of downregulated let7-b in prostate cancer^43^. The range obtained from the prostate cancer sample here is in excellent agreement with relative expression ratios of miR-miR-141 and miR-375 reported at 1.9- and 2.6-fold, respectively, in a study including 20 Prostate Cancer patients and 8 healthy individuals^41^. The observation of less than a factor of 3-fold difference between cancer samples and healthy ones suggests that these miRNAs need to be measured accurately and correlated to the serum RNA (see later).

### Multiplexing probes to assess combined copy number of three miRNAs

To show multiplexing of probes and internally validate the levels of miR-21, miR-375 and miR-141 determined in healthy and cancer samples we designed experiments that contained all three probes. In this assay multiplexing works only for miRNAs that exhibit comparable levels because the nanopores cannot discriminate between one probe or another. Optimization of the ONT flow cells and/or the *OsBp_detect* algorithm towards probe discrimination is possible. Earlier studies have shown that (*I*_*r*_*/I*_*o*_)_max_ varies as a function of the number of OsBp moieties with less OsBp resulting to a later (*I*_*r*_*/I*_*o*_)_max_ while more OsBp yields an earlier (*I*_*r*_*/I*_*o*_)_max_. Iin the absence of optimization to yield probe discrimination, multiplexing may be a useful approach to test simultaneously the levels of two proven cancer biomarkers, such as miR-375 and miR-141 which measure 4,620 and 3,048 copies, respectively. Multiplexing could be also used to test the level of a validated miRNA biomarker for a specific disease in combination with miR-21 that appears to be overexpressed in many diseases^44^.

We chose 3 probes to target miR-21, miR-375 and miR-141 in H6914 and considered the copy number of each target obtained from the corresponding silencing experiment as low level (LOW HL), and the copy number of each target obtained from the corresponding detection experiment as high level (HIGH HL). Probe content was matched to its target. Experiments were conducted with the small RNA fraction and repeated with total RNA. Three different flow cells were explored at different stages of their lifespan. It was determined that the level of the combined three miRNAs is more than the LOW HL and less than the HIGH HL. The same result was obtained from either the small RNA or the total RNA fractions (Fig. S6 in Supplementary). This set of experiments validates the use of the Monarch kit for RNA purification, the use of the total RNA fraction as test article, and multiplexing probes to target miRNAs with comparable counts.

### Correlating miR-15b copies with total RNA in the sample

The observed 2 to 3-fold overexpression of cancer biomarkers in Table 2 appears too small of an effect to judge someone’s health status. RNA content in serum may be partially responsible for a biomarker’s level. Total RNA isolated from serum measures 2 to 20 ng/μL, as determined using a nanodrop spectrophotometer. Instead of a nanodrop, we chose a Capillary electrophoresis (CE) Agilent instrument (see Methods and Fig. S7 in the Supplementary). Table 3 lists the accumulated data of miR-15b copies as a function of total RNA (Fig. 6). Serum samples from healthy and cancer patients that exhibit total RNA below 5ng/μL were not subject to nanopore experiments, because measurements below 5ng/μL RNA carry a large standard error.

**Figure 6.**
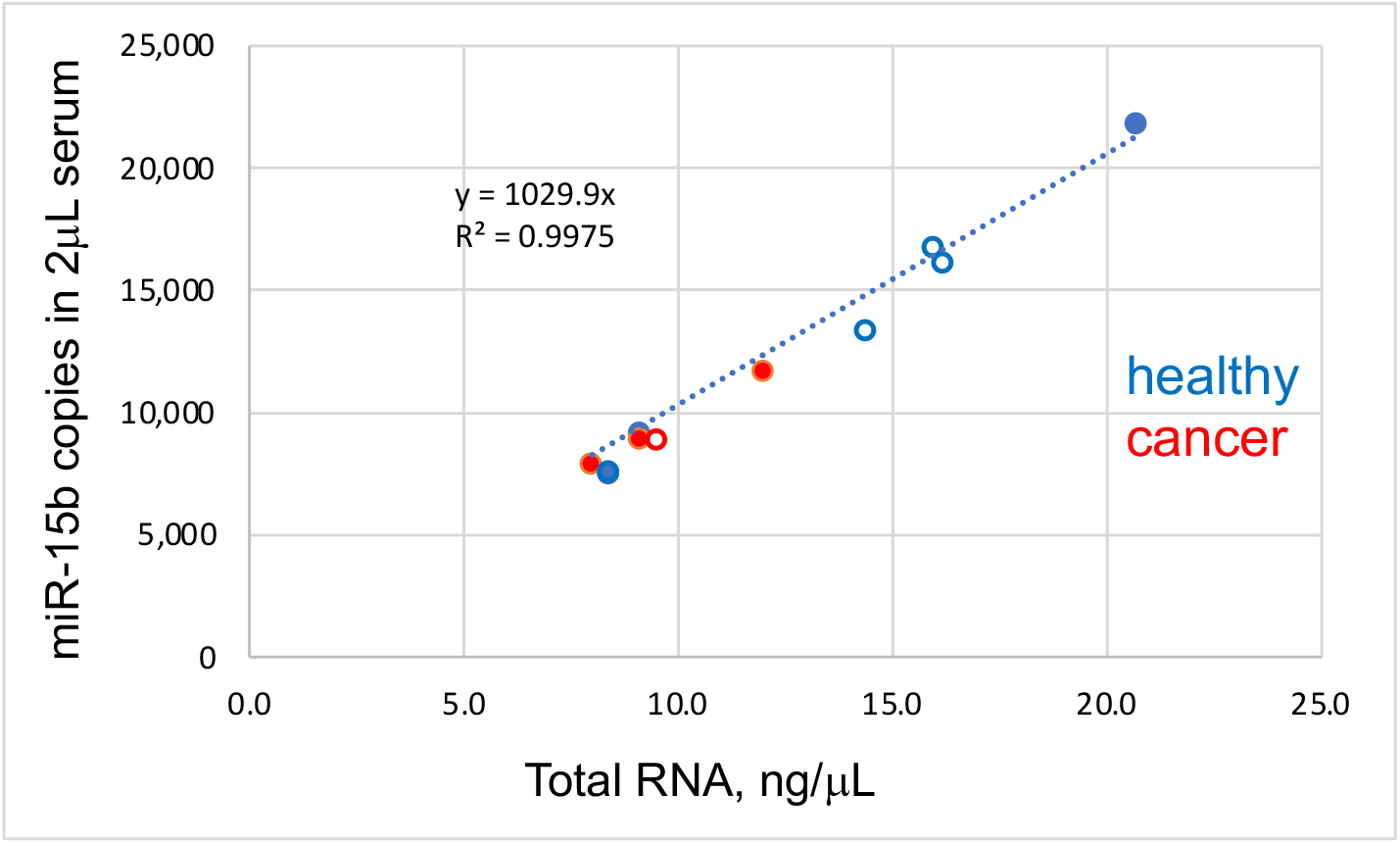
(see Table 3): Copies of miR-15b per μL of total or small RNA (2uL serum) as a function of total RNA in ng/μL isolated from serum. Total RNA was determined by CE using an mRNA standard (Methods). Concentration of the mRNA standard and some of the serum samples were confirmed by a nanodrop spectrophotometer. Copies of miR-15b were determined using the MinION (Table S2 in Supplementary). Total RNA or small RNA was isolated from serum using the Monarch RNA purification kit (NEB, see Methods). Total RNA and Small RNA are equivalent, but 2-fold concentrated over serum. Copy numbers of miR-15b determined from H4522, either as total RNA or small RNA are artificially identical, because the experiments were identical by design. These data suggest that (i) total RNA or small RNA obtained using the Monarch kit are reliable sources for miR-15b quantification and (ii) miR-15b copies are comparable among healthy and cancer patients, after normalization against the RNA serum content.

Experiments in Fig. 6 were conducted using small RNA or total RNA fraction, RNA obtained from two lots of H6914, as well as from healthy individuals or cancer patients. The perfectly linear correlation in Fig. 6, confirms earlier reports that miR-15b copies are comparable among healthy and cancer patients^6^, and illustrates that: (a) total RNA and small RNA fractions obtained from the Monarch purification kit exhibit comparable amounts for miR-15b, and likely for other miRNAs. (b) The range of 7,500 to 22,000 miR-15b copies correlates linearly with total RNA in 1μL of total or small RNA or 2μL serum (see Methods) and the correlation goes through the intercept 0,0, as it should. Presumably this correlation can be extended at both ends and future work may test for such correlation with other miRNAs.

Studies to quantify miRNAs are concerned with specificity and single nt discrimination. Many technologies depend on hybridization efficiency between a probe and the target miRNA. Since the concentration of miRNAs in serum are typically in the fM range, our experiments are designed to target the fM range. We have optimized the conditions for probe and target hybridization (overnight at -20°C^45^, ahead of buffer addition, see Methods) to assure complete hybridization. With other technologies discrimination is achieved by improving the hybridization efficiency. For example, it was reported that including a locked nucleic acid (LNA)-mediated PCR step upstream of electrochemical detection improved the limit of detection (LOD) by 2 orders of magnitude, down to an ultralow-level of 5 copies per μL^46^. Since hybridization efficiency is already optimized in this study, to achieve specificity we envision lessening the hybridization efficiency by selectively incorporating two consecutive T(OsBp) moieties within the complementary sequence to partially distort single strand configuration and enable specificity between miRNAs that differ by one or two nucleotides.

The need for novel PCR-free technologies to measure miRNAs has led to several innovations. Direct kinetic fingerprinting for high-accuracy counting of diverse disease biomarkers (SiMREPS) uses single-molecule fluorescence microscopy and reports proof-of-principle studies with LOD at 1 fM and better than 500-fold selectivity for single-nt polymorphisms^47,48^. A recent application of the MinION sequencing platform uses barcoded probes for multiple analytes including miRNAs with a reported miRNA LOD at 50pM^49^. Electro-optical nanopores and size-coded molecular probes were exploited for extraction- and amplifcation-free multiplexed miRNA detection from human serum^32,43^. This experimental nanopore platform demonstrated the possibility to classify prostate cancer stages by simultaneously profiling a panel of 3 miRNAs and obtained LOD at about 100 fM for miR-375-3p and miR-141-3p^43^. The 100 fM LOD is over 20-fold higher compared to our current LOQ at 5 fM miR-141, the lowest miRNA copy number determined here (3.048 in Table 2). If our experiments identify a miRNA that exhibits fewer than 3000 copies, detectability can be improved by 4-fold and approach 1fM. This can be accomplished using a 2-fold concentration in the RNA isolation step and reducing probe molecules by 2-fold. In contrast to the MinION-based sensing platform described here, many promising technologies are still at the experimental stage and require expensive infrastructure and skilled personnel, which prevents their use by less experienced end-users or technicians at a POC facility.

## Conclusions

In summary, we developed and validated a low-throughput, nanopore array-based assay using a commercially available nanopore platform. We measured miRNA copies from serum as low as 5fM with better than 85% confidence. This was accomplished by exploiting an RNA purification/isolation kit, the MinION nanopore device with consumable flow cells, and a proprietary probe complementary to each miRNA target. The technology uses the MinION, but instead of sequencing, conducts 45 min long single-molecule ion-conductance experiments at -180mV. The raw *i-t* data are analyzed by *OsBp_detect*, a publicly available software package developed specifically for this application^34^. The probe molecules are preferentially detected over intact RNA and traverse the MinION nanopores quantitatively. The probes can be either DNA, RNA, or 2’-OMe-oligos. The design of the probes includes a sequence complementary to the target while replacing the complementary Ts by Us, adding extra Ts at one end, and A-tails at both ends. The Ts are osmylated by 30 min incubation with an inexpensive reagent, OsBp, at room temperature. The conjugated T-OsBp moieties are responsible for the remarkable discrimination between intact oligos and probes.

What stands out compared to other miRNA quantification methods is that this technology is PCR-free, simple and ready-to-use, and the probes are easy to design, manufacture and are chemically stable. Most importantly, miRNA copies are measured directly and not relative to another miRNA, i.e., there is no normalization employed. This assay was validated in the range 3,000 to 100,000 miRNA copies 1 μL serum, exhibits better than 85% confidence across this concentration range, and currently showcases a LOQ at 5 fM (3,048 miR-141-3p copies in 1μL serum from H6914). The data exhibit a linear correlation between miR-15b copies and ng/μL RNA, as obtained by total RNA isolation from serum and quantified by traditional analytical instruments. The correlation included miR-15b determinations in samples from the combined serum of healthy individuals H6914, from healthy individuals, and from cancer patients with pancreatic, prostate or breast cancer. Linear relationships between miRNA copies and isolated RNA may apply to other miRNAs besides miR-15b, and likely differentiate healthy from diseased samples. The exceptional agreement of relative abundance for miR-21/miR-15b obtained from H6914 and from a healthy individual H1 measuring 0.59 and 0.61, respectively, validates this approach. Due to its unprecedented precision, this assay may be used as a QC test to confirm miRNA copies obtained by high-throughput technologies. It could be exploited as an LDT to diagnose cancer or other diseases from a few validated biomarkers, perhaps at the earliest stage possible when miRNAs become dysregulated. If miRNAs prove to be the “tiny regulators” that Victor Ambros proposed them to be^3^, then this LDT and others, based on the principles described above, may be implemented at POC facilities, part of regular check-ups for adults across the globe.

## Methods

### Serum samples, Oligos, and other Reagents

Human Serum from human male AB plasma (H6914, 1^st^ lot SLCH8785, 2^nd^ lot SLCJ3635) and (H4522, lot SLCK9619) were purchased from Sigma-Aldrich. Serum samples from healthy individuals and cancer patients, preferably pretreatment, were purchased from the Discovery Life Sciences blood bank (DLS, Table S1, Supplementary). For the isolation of RNA from serum we used the Monarch T2010S kit (older lot 10075450, newer lot 10141109). Custom-made RNA oligos, deoxyoligos and 2’-OMe-oligos, typically synthesized at the 0.2 μM scale, and purified by the manufacturer were purchased from Dharmacon (Horizon Discovery Group), Integrated DNA Technologies (IDT) and Millipore/Sigma-Aldrich, respectively. Oligos (sequences in Table 1) were diluted with Ambion Nuclease-free water, not DEPC treated, typically to 100 or 200 μM stock solutions and stored at −20°C. Oligos purity was tested by HPLC in-house and found to be >85%^50^. The only ONT kit required for the experiments reported here is kit SK1581018XL, ONT flush buffer or ONT buffer. ONT buffer is proprietary, provides the necessary electrolytes and must represent over 85% of the 75μL sample volume. Buffer, DNase-free and RNAse-free, TRIS.HCl 1.0 M pH 8.0 Ultrapure was purchased from Invitrogen, NaCl crystalline ACS min 99.0% from Alfa Aesar, and distilled water from Alhambra or Arrowhead. These three materials were used to prepare the HPLC mobile phase to test oligo purity, determine probe content, extent of osmylation, and confirm probe/target hybridization. A 4% aqueous osmium tetroxide solution (0.1575 M OsO_4_ in ampules at 2 mL each) was purchased from Electron Microscopy Sciences, and 2,2’-Bipyridine 99+% (bipy) from Acros Organics. LoBind Eppendorf test tubes (1.5 mL) were used for 1/100 probe dilutions to yield probes at 15 to 30fM concentration. Samples of probe and serum RNA were prepared in 0.5 mL RNase-, DNase-free, sterile test tubes.

### Manufacturing of OsBp-nucleic acids has been reported^28-30^ and is summarized here

OsBp reagent is prepared by weighing the equivalent of 15.7mM of 2,2’-bipyridine or bipy (49.2mg) in a 20mL scintillation vial, adding 18mL of water and stirring at room temperature until bipy dissolves. The full content of a 2mL 4% OsO_4_ solution, supplied in an ampule, is transferred to the vial using a glass pipette. This mixture is an aqueous 20mL 15.75 mM OsBp (0.4%) stock solution, equimolar in OsO_4_ and bipy. OsO_4_ is a hazardous material^51^. Care should be taken that this preparation, as well as any other work using OsBp is conducted in stoppered glass vials in a well-ventilated area. Leftover solutions of OsO_4_ and/or OsBp may be mixed with corn oil to neutralize unreacted OsO_4_ and properly discarded^51^. The freshly prepared OsBp stock solution is dispensed in HPLC vials and kept at -20°C. Each vial can be stored at 4°C and used for a week without loss of potency; typical pipette tips can be used for manufacturing of OsBp-labeled nucleic acids. OsBp stock solutions should be validated before first use by running a known reaction. For osmylation reactions a 20-fold excess of OsBp over the reactive pyrimidine in monomer equivalents is required to ensure pseudo-first order kinetics and yield preferential labeling of T over the other pyrimidines. Quenching of the osmylation reaction occurs upon purification. Purification from excess OsBp (twice) was done with spin columns (TC-100 FC from TrimGen Corporation) 4 min at 5,000 rpm. The flow-through solution is the purified osmylated oligo. The conjugated pyridine-OsBp moiety is unreactive, osmylated nucleic acids are stable and may be stored in 1.5 mL microcentrifuge tubes at -20ºC for a year.

### Probe Optimization for MinION detection

The probes are derived from short oligos, DNA or RNA, as is, or with some or all 2’-OMe-moieties. They are around 38nt long, each probe being unique to its target (Table 1). Earlier work indicated that any oligo that carries one or more OsBp moieties traverses and can be detected by α-hemolysin^28^, by sufficiently narrow solid state nanopores^27^, as well as by the proprietary nanopore of the MinION flow cells^29^. To quantify miRNAs from serum, it is imperative that the assay is relatively quick, let us say about an hour, and that the probe is quantitatively detected within this time frame. Earlier experiments using probes with 2 or 3 OsBp moieties indicated that only 10 to 20% of the probe molecules yielded events within an hour. Experiments with probes that contain 4 to 8 OsBp moieties but no A-tails at the 3’-end or 5’-end also exhibited poor nanopore detection. The optimized probe design is comprised of a sequence complementary to its target but extended at the 5’-end or at the 3’-end with 4 to 5 adjacent Ts (one end only) and flanked by up to 5 As at either end (see Table 1). The As facilitate oligo entering the nanopores. The adjacent Ts are conjugated with OsBp and responsible for quantitative detectability. Within the probe sequence complementary to the target, Ts are replaced by U, 2’-OMe-U or dU, to minimize OsBp-labeling because the kinetics of Uridine (U), U-derivatives and of Cytosine (C) osmylation, are substantially slower compared to T^28, 30,39^.

### Occasional single OsBp moiety present in the complementary portion of the probe does not prevent efficient hybridization

In Table S2 under flow cell FAT43745 (Supplementary) there are experiments using a probe preparation for miR-21 that carries 6.1 OsBp moieties and a probe preparation for miR-141 that carries 6.9 OsBp moieties. Experiments with these probes yielded both detection and silencing, clearly suggesting that up to one OsBp moiety per 10 nt (6.9 OsBp is about 2 OsBp per 22nt miRNA sequence) doesn’t distort the configuration of the ss molecule and therefore doesn’t prevent hybridization with the target. Due to the randomness of the osmylation process, the 2 OsBp moieties in miR-141 are expected to be primarily single T-OsBp and not adjacent. Using probe miR15bT5 and conditions for partial T-osmylation, we discovered that RNA probes with more than two adjacent OsBp at the 3’end do not cross the MinION pores, while two adjacent OsBps at the 3’end yield quantitative detection within 45 to 90 min (see Table S2 in the Supplementary section under probe miR15bT5 (1.5 OsBp)). DNA probes require four or five adjacent OsBps for quantitative detection. This observation can be rationalized by the aqueous solvation of the 3’-OH in RNA which requires more nanopore space for RNA compared to DNA and the fact that the conditions of the nanopore experiment may reduce the bulk of water around a nucleic acid without removing the first aqueous solvation shell (see discussion in reference 29).

*(Pyrimidine)OsBp is a chromophore and absorbs* in the range of 300nm to 320nm^30,39^, where nucleic acids exhibit negligible absorbance. We exploited this observation and showed that extent of osmylation can be measured using the equation: Absorbance at 312nm/Absorbance at 272nm or R(312/272) = 2x(No of osmylated pyrimidines/total nt)^30,39^. R is the ratio of the corresponding HPLC peaks, regardless if these peaks are sharp, broad or multiple. The wavelengths 312 nm and 272 nm were chosen to maximize the effect and to equalize contributions by different pyrimidines^28-30,39^. The number obtained from the equation refers to OsBp moieties, on average. Because a molecule carries an integer number of OsBp moieties, some molecules will have fewer, and some molecules will have more than the determined average value. Osmylation occurs randomly, including adjacent pyrimidines, but depends on the relative reactivity of OsBp for a pyrimidine; trends are comparable in deoxy- and ribodeoxy-nucleic acids^28,29^. Because of the markedly higher reactivity of OsBp towards T, compared to U and C, manufacturing under our conditions (2.62mM OsBp (1:1) with 30 min incubation at room temperature), yields over 90% T(OsBp) and less than two (U(OsBp) + C(OsBp)) within the 22nt sequence of the probe complementary to the target miRNA. We use HPLC analysis to obtain probe concentration (content) after purification by using the intact oligo as a standard, because absorbance of the osmylated oligo is practically the same as the precursor intact oligo^39^. HPLC is useful as it provides evidence for quantitative depletion of the OsBp reagent. Alternatively, a nanodrop spectrophotometer can be used for content and extent of osmylation.

### HPLC methods for probe purity, content, osmylation extent, and Hybridization with target miRNA

The HPLC method to assess oligo purity, content, and osmylation extent was developed earlier^50^. A comparable method was developed and used earlier to assess hybridization between a certain miRNA and its osmylated probe in 85% ONT buffer^30^. Figure S1 (Supplementary Section) illustrates HPLC evidence for hybridization obtained for let7-b and probe let7-b(T5) to suggest that the adjacent T(OsBp) moieties and A-tails do not interfere with hybridization in 85% ONT buffer at the 5 μM level. Because probe and RNA are mixed at single digit fM concentrations there was concern regarding incomplete hybridization. Incubation of the mixture for couple of hours at room temperature was found inadequate by nanopore testing. The condition that proved optimal was overnight storage at - 20°C, a condition known to yield an impressive concentration of the organic components upon freezing^45^. ONT buffer was added to the probe/RNA sample right before the nanopore experiment. HPLC analysis was done using an Agilent 1100/1200 LC HPLC equipped with a binary pump, Diode Array Detector (DAD), a 1290 Infinity Autosampler/Thermostat, and Chemstation software Rev.B.04.01 SP1 for data acquisition and processing. IEX HPLC column DNAPac PA200 from ThermoFisher Scientific (Dionex) in a 4X250mm, or in a 4X50mm configuration were used. HPLC analyses were conducted automatically using a thermostatted autosampler at 15°C. HPLC peaks were detected and identified using a diode array detector (DAD) in the UV–vis region 210–450 nm. The chromatograms were recorded at 215, 260, 280, 272, and 312nm. Samples were prepared with RNase free water, and buffers by commercial distilled water. The HPLC method uses mobile phase A (MPA) aqueous 25mM TRIS.HCl pH 8 buffer, mobile phase B (MPB) aqueous 1.5M NaCl in a 25mM TRIS.HCl pH 8 buffer, a gradient from 10% MPB to 60% MPB in 20min and an additional 10min for wash and equilibration to initial conditions, i.e., 90% MPA. Column temperature was set at 35ºC and flow rate was 1mL/min or 1.3mL/min depending on column configuration (see above). ONT buffer is proprietary material. Whether the ONT component observed by HPLC in the ONT buffer (Fig. S1) and the ONT component seen in the *i-t* traces of the nanopore experiments (Fig. S2) are the same material is unclear.

### Isolation of total RNA and small RNA from serum using the Monarch NEB kit T2010S

We revised slightly the protocols of the manufacturer to isolate total RNA (A) and small RNA (B). (A) Manufacturer’s instructions were used to reconstitute DNase I, Proteinase K add to add 95% ethanol to Wash buffer. All spinning was done here at 15,000 rpm and for 1 min. Protocol: Thaw blood/serum, take 0.2 mL and add 0.2 mL of DNA/RNA Protection Reagent. Vortex briefly, add 10 μL of reconstituted Proteinase K, vortex and incubate at room temperature for 30 min.

Add 0.4 mL of isopropanol, vortex, and transfer mixture to an RNA purification column (dark blue n) fitted with a collection tube. Spin for 1 min. Discard flow-through. Add 0.5 mL RNA Wash Buffer and spin for 1 min. Discard flow-through. In an RNase-free microfuge tube combine 5 μL reconstituted DNase I with 75 μL DNase I Reaction Buffer and pipet directly to the top of the column matrix. Incubate for 15 minutes at room temperature. Add 0.5 mL RNA Priming Buffer and spin for 1 min. Discard flow-through. Add 0.5 mL RNA Wash Buffer and spin for 1 min. Discard flow-through. Add another 0.5 mL RNA Wash Buffer and spin for 3 min. Discard flow-through. Transfer column to an RNase-free microfuge tube. Use care to ensure the tip of the column does not contact the flow-through. Add 100 μL Nuclease-free Water directly to the center of column matrix and spin for 1 min. The flow-through is 0.1 mL of total RNA isolated from the 0.2 mL blood/serum.

(B We isolated the small RNA fraction (less than 200nt) from 50 μL of the above 0.1mL total RNA. Protocol: Prepare an equal mixture of RNA Lysis Buffer and 95% ethanol; add 100 μL to the 50 μL total RNA sample. Mix by pipetting. Transfer it into an RNA Purification Column (dark blue) fitted with a collection tube and spin for 1 min at 15,000 rpm. Save the flow-through as RNAs > 200 nt bind to the column, whereas RNAs < 200 nt partition into the flow-through. Add 0.5 mL RNA Priming Buffer and spin for 1 min. Discard flow-through. Add 0.5 mL RNA Wash Buffer and spin for 1 min. Discard flow-through. Add another 0.5 mL. RNA Wash Buffer and spin for 3 min. Discard flow-through. Transfer column to an RNase-free microfuge tube. Use care to ensure the tip of the column does not contact the flow-through. Add 50 μL Nuclease-free Water directly to the center of column matrix and spin for 1 min. The flow-through is 50 μL of small RNA.

*Capillary Electrophoresis (CE) analyses* were conducted with an Agilent G1600 Capillary Electrophoresis instrument using electrokinetic injection and equipped with DAD and Chemstation software Rev.B.04.03 for data acquisition and processing. The CE was used in conjunction with a circulating bath to control the autosampler’s temperature at 15 °C. The capillary’s temperature was controlled by the instrument’s software at 25°C. Capillary zone electrophoresis (CZE) analyses (see Fig. S7 in Supplementary) were conducted with an untreated fused silica capillary (50 μm × 40 cm) with an extended light path purchased from Agilent Technologies in pH 9.3 and with 50 mM sodium tetraborate buffer using a CE method at 20kV and 24s injection time.

### Single molecule ion-channel conductance experiments on the MinION (MinION Mk1B platform)

To use the ONT platform one must register with the company and download the software MinKNOW to a computer/laptop (specs are provided by ONT). All the functions necessary to test the hardware, the flow cell, and run the experiments are done via the MinKNOW software tool. Raw data files are acquired in *fast-5* format at about 3.5 GB and analyzed by the *OsBp_detect* software^34^. MinKNOW allows for monitoring in real time any chosen channel, and *fast-5* files can be directly visualized in MatLab (from Mathworks) 2D format once the experiment is completed. The heart of the MinION platform is the consumable flow cell; both R9 and R10 versions are used successfully. The Flongle, also supplied by ONT, was found less useful for this application because it is not recommended for multiple experiments. With the MinION flow cell, up to twenty 45-min nanopore experiments can be conducted following the protocol below.

The MinION device loaded with the flow cell/sample is connected to a computer/laptop carrying the ONT software, MinKNOW. The latter runs the experiment and acquires the data (*i-t*). *OsBp_detect* is also downloaded to a laptop/computer and used to analyze the *fast-5* files and report events per channel per threshold values set by the user.

### Running a nanopore experiment with Yenos probes

The sample is loaded on the flow cell that sits on the MinION device. The experiment is run under “start sequencing” mode using the MinKNOW software (we used here vs. 21.11.7). Direct RNA Sequencing Kit (SQK-RNA002) is selected just to run the experiment. Run Length (45 min), Bias Voltage (−180 mV) are selected, Basecalling is disabled and Output bulk file - Raw (1-512) enabled. Output location is /Library/MinKNOW/data/., and Output format is *fast-5*. All the experiments reported here run for 45 min at -180mV. Higher voltage was tested and found to lead to faster pore inactivation. A 45-min experiment produces a 3.3 GB data file, which can be analyzed using our *OsBp_detect* algorithm within minutes^34^. While alternative parameters were tested, all the experiments reported here were analyzed using threshold parameters as listed below (see Protocol 8. In Methods). A state-of-the-art laptop/computer may require less than 5 minutes for the *OsBp_detect* analysis of the 3.3GB fast-5 file and produce a file, in tsv format, which can be opened via Microsoft Excel and saved as such. In the Excel spreadsheet, the data are grouped in the form of a histogram with 0.05 bins, from 0.05 to 0.55, and plotted (see Figures). Number of events per channel from *OsBp_detect* analysis was compared to the actual *i-t* trace of the specific channel by MatLab visualization, and this algorithm, 2^nd^ revision, was repeatedly shown to work perfectly.

### Protocol/Workflow for miRNA quantification

1. Use NEB kit T2010S and isolate RNA from the serum/blood sample (see our protocol above).
2. Obtain flow cells R9 or R10 from ONT and test for acceptable number of pores.
3. Open both the priming port and the sample port and load using a 200 μL pipette tip 0.6 to 1.0 mL of ONT buffer, filtered via Sterlitech aluminum oxide 0.02-micron filters. Immediately remove excess via the priming port using a 1000 μL pipette tip. Make sure that no air bubble enters the nanopore array, as this will inactivate many nanopores. Air bubbles inside the array are impossible to remove.
4. Obtain tested Yenos osmylated probes and dilute with Ambion water to about 15 to 25 fM.
5. Mix a few μL RNA with a few μL of a Yenos probe and keep at -20°C overnight to assure successful hybridization^41^.
6. Add 75 μL of filtered ONT buffer to the sample (5.) and run the MinION experiment for 45 min at -180mV.
7. Let the flow cell rest for 15 min at room temperature and run another 45 min at -180mV without opening the flow cell (nonew). Store the flow cell at 4°C between experiments and conduct no more than two 45-min experiments at a time and no more than four 45 min experiments per day with the same flow cell. The first experiment to run on a new flow cell, after completing step 3. is a buffer test to establish that within a 45min run a R9 cell exhibits less than 200,000 events and a R10 cell less than 300,000 events. Every experiment produces automatically a *fast-5* file with the raw data.
8. Analyze the *fast-5* file using the *OsBp_detect* algorithm with thresholds: (i) event duration (in tps): 4-1200, (ii) lowest *I*_*r*_*/I*_*o*_ <0.55, (iii) all *I*_*r*_*/I*_*o*_ <0.6, channels 1-512. Discrimination based on relative residual ion current (*I*_*r*_*/I*_*o*_) is better compared to discrimination based on residence time τ (appearing skinny or fat in Fig. S2 (Supplementary)) because τ distribution is broader compared to *I*_*r*_*/I*_*o*_ distribution. The histogram data may or may not be graphed. The algorithm-selected events are to be added together, **Total Events**, the late (*I*_*r*_*/I*_*o*_)_max_ and early (*I*_*r*_*/I*_*o*_)_max_ identified, and their **Ratio** calculated. These values represent the criteria by which an experiment is judged as detection or silencing experiment (see under 9. and discussion in text). Future *OsBp_detect* versions may incorporate this analysis, compare experiments, and yield conclusions.
9. To determine if an experiment (E) that contains probe and RNA, is a detection or a silencing one: Compare experiments conducted on the same flow cell and done sequentially with probe at an approximate 50,000 load. As control (C) use the 2^nd^ run (nonew) of an experiment such as a buffer test, a probe in buffer, or RNA in buffer. Compare Total events and the Ratio (see under 8.) between E and C. A buffer test and an RNA in buffer experiment is expected to yield fewer events compared to a probe in buffer experiment. A probe in buffer experiment and a detection experiment may exhibit comparable events and, at least, 2-fold more compared to a buffer test or a silencing experiment. If Total Events are comparable between E and C, then an increased Ratio indicates silencing, whereas a decreased Ratio indicates detection.
10. Determine copies of a target miRNA as follows: Select two experiments that differ by about 20% in small RNA load, one exhibiting detection and the other silencing, calculate for each experiment the copies of target miRNA = probe molecules/(RNA in μL) and take the average, which will approximate miRNA copies per 1μL of the RNA sample with probe molecules = (probe μL) x (probe concentration in fM) x 600.

## Data availability

The data generated during this study are included in this published article and/or in the Supplementary Section. Raw data (*fast-5* format at 3.4 GB each) may be obtained from AK.

## Supplementary Information

see separate file, online.

## Acknowledgements

The work was supported by private funding and enabled using the *OsBp-detect* algorithm developed specifically for this application by Dr. Albert Kang. We are immensely gratefully for his contribution and his support in revising it, as needed. We are thankful to Dr. William Jack from New England BioLabs for recommending the Monarch RNA purification kit. We also thank Professor Miten Jain from Northeastern University, for sharing with us the algorithm to visualize the *i-t* trace of each channel of the *fast-5* file using MatLab from Mathworks.

## Additional Information

### Competing Interests

Anastassia Kanavarioti is the Founder and Director of Yenos Analytical LLC, a company delivering custom analytical solutions for native, synthetic, or transcribed nucleic acids, and engaged in the development and manufacturing of labeled/tagged nucleic acids for use in conjunction with nanopore detection and nucleic acid quantification including miRNAs ([Nanopore platform for DNA/RNA oligo detection using an osmium tagged complementary probe](https://patents.justia.com/patent/11111527), Patent No: US 11,111,527 B1, issued 9-7-2021 and US 11,427,859, issued 9-30-2022). There is no competing interest or relationship (commercial, financial, or non-financial) between the two companies, namely Oxford Nanopore Technologies, and Yenos Analytical LLC and their employees.

