## Supplementary for "Femtomolar-level PCR-free quantification of microRNA cancer biomarkers in serum"

**Anastassia Kanavarioti**

Yenos Analytical LLC, 4659 Golden Foothill Pkwy, Suite 101, El Dorado Hills, CA 95672, USA

#### Supplementary Section

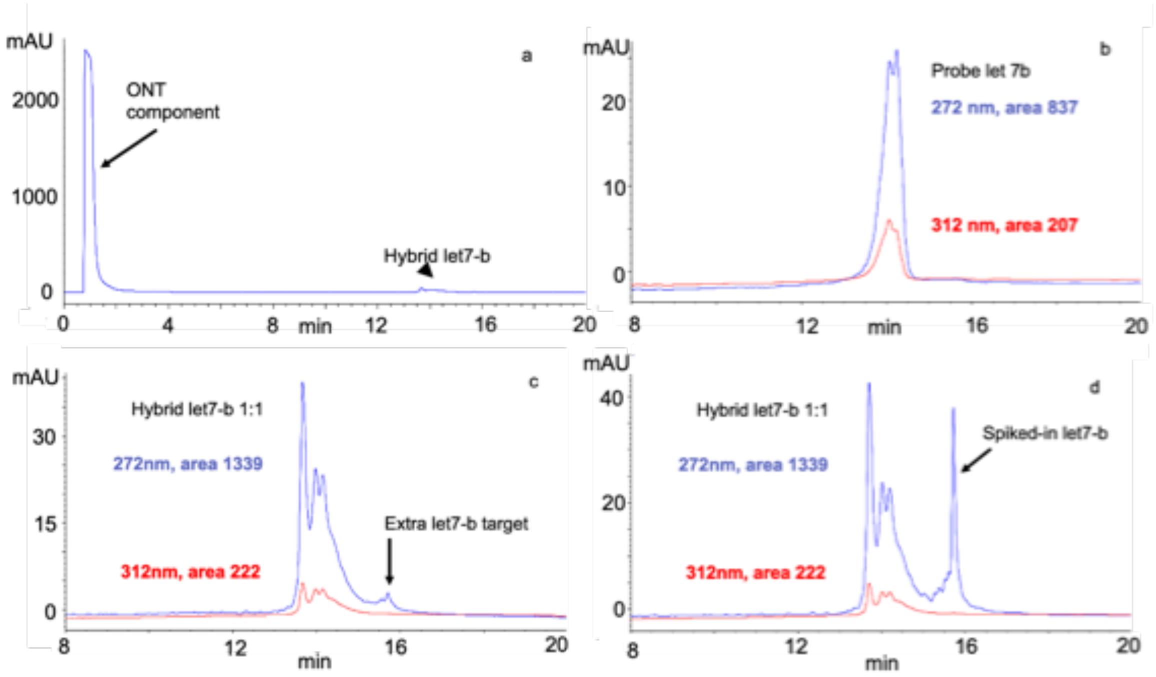

**Figure S1: HPLC profiles of samples to show 1:1 hybridization between let7-b and probe let7bT5 (Table 1 and Methods under HPLC).**

**(a)** Sample with 20 pmoles of the hybrid formed in a 1:1 solution of let7-b and probe let7bT5 (4.4 OsBp) in 85% ONT buffer. The huge peak at void volume is identified here as an ONT component with chromatographic, electrophoretic and spectroscopic characteristics comparable to Adenosine triphosphate (ATP). The hybrid elutes at about 14 min and is barely detectable due to the excess of the ONT component found in the ONT buffer. This chromatography resolves ONT component from probe, target or hybrid for most target/probe sets tested. **(b)** Sample with 20 pmoles of Probe let7-bT5. The value R=(HPLC area at 312nm)/(HPLC area at 272nm)=207/837=0.247, yields OsBp=4.4 moieties on average (from 0.247x36/2) and implies extensive osmylation of the 5 Ts, but negligible osmylation elsewhere due to the high selectivity of T over the other pyrimidines under our osmylation conditions[29,30,39]. **(c)** Same sample as in (a), magnification to show the hybrid. The ratio R=222/1339=0.16 is practically equal to R(probe)/2, consistent with a 1:1 hybrid between the osmylated probe and the intact target. A small peak eluting later is attributed to unhybridized let7-b. **(d)** HPLC profile of the hybrid sample (c) after spiking-in intact let7-b to confirm the identity of the small peak observed under (c).

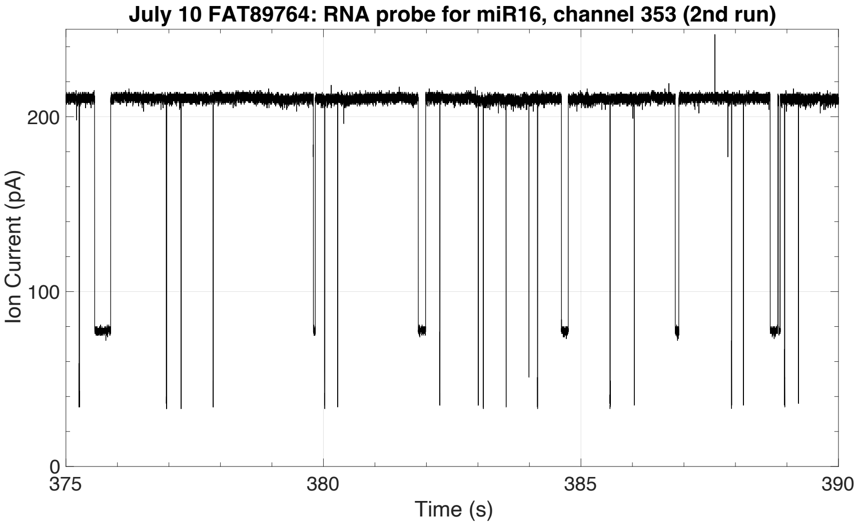

**Figure S2: Trace *i-t* via MatLab visualization of the fast-5 file. It illustrates visual discrimination of events attributed to a Yenos' probe (long and skinny) and the ONT component(s) (short and fat).** Events attributed to the ONT component(s), present in the solution that the flow cell is supplied with, exhibit $I_r/I_o \cong 0.33$. Events attributed to Yenos' probes exhibit here $I_r/I_o \cong 0.15$. This figure was obtained from an experiment with probe for miR-16 (RNA rT4probe16, see Table 1) at 45,000 counts in buffer.Please note that while the MinION flow cells have 2048 nanopores, there are only 512 channels to report *i-t*.

Some observations made with the ONT nanopore array may apply to future nanopore arrays. Even though the 75μL sample is a homogeneous solution, the data/events obtained from each working pore indicate inhomogeneity. This inhomogeneity is discovered by looking at the number of events per channel, determining an average and finding that among the working pores there is typically a 5% of "outliers". We described these outliers as pores that detect/report 5 to 25-fold more events compared to the average number of events detected in the remaining 95% of pores[29,30]. Repeated investigations into this phenomenon lead us to suggest that variations of the applied voltage, malfunctioning pores, or the existence of extra sensitive pore protein variants within the batch cannot explain the presence of outliers.

An example of an outlier is seen in this figure: Fig. S2 includes 24 events, by eye, within 15 s of run, 863 events as reported using *OsBp_detect* for this channel and a total of 55, 969 events from the experiment with 353 working nanopores out of 874 available ones, as reported by MINKNOW at the beginning of the experiment. The average number of events per channel is 158 from 55,969/353, and this value is confirmed by eye from the *OsBp_detect* excel file. The 863 events are a 5.5-fold higher than the average and define this channel as an outlier. The fact that this channel reports both events from both materials is a common occurrence, whereas outliers as mentioned earlier are in the range of 5% of reporting channels.

An outlier differs from a malfunctioning pore in that the outlier channel reports 5 to 25-fold more events, and typically an average number of events in the next run (nonew), most likely because the inhomogeneity that produced the outlier has moved elsewhere or disappeared, while the voltage is turned off. On the contrary, malfunctioning pores that exhibit an abnormally large number of events, are rare at about 1 such pore per 10 flow cells, and persist from one experiment to the next, including new samples, until they eventually become inactive.

With respect to what creates the described inhomogeneity in the first place, we offer the following rationalization: Negatively charged molecules need to travel a distance before reaching a pore and traverse. This takes time and because of the extra low (fM) concentration experiments exhibit around 100 to 300 events per channel or 50,000 to 150,000 total events per 45min long experiment (our parameters, see Protocol 8. in Methods). We propose that inhomogeneity happens when a small number of charged species happens to be already at the proximity of a pore. This small number of molecules traverses at the onset of the applied voltage and creates a “void” which in turn creates a force, in addition to the voltage, and yields a flux of charged molecules to fill the void and traverse this pore. We envision this phenomenon as a river of charged molecules that rushes towards the specific pore creating the “outlier”. Outliers are observed with flow cells that have most of the 512 pores active, as well as with flow cells that have only very few active pores left. It explains how the number of pores of a flow cell may decrease substantially but the number of observed events doesn’t follow proportionally. We have analyzed data by either including or excluding the outliers, and the conclusion of the experiments didn’t change[30]. In this study no channels were excluded.

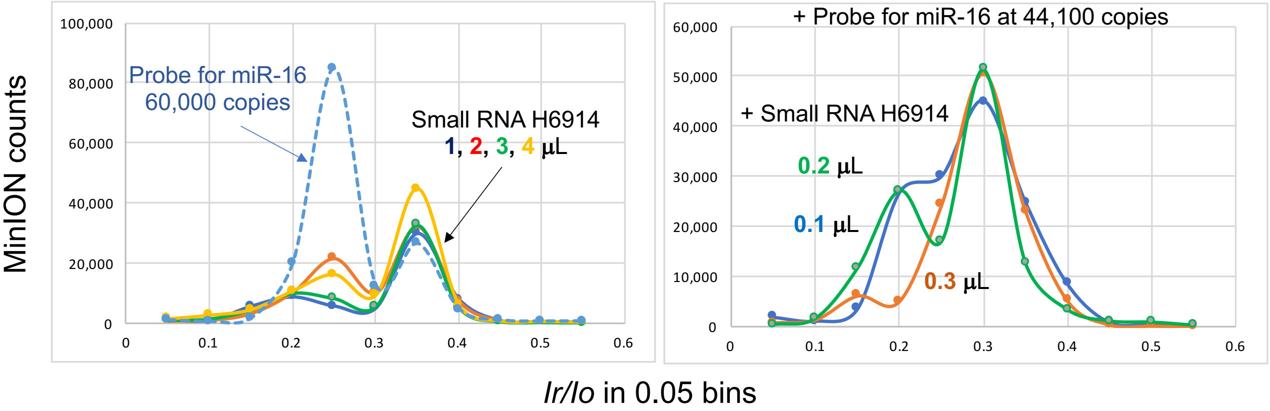

**Figure S3a: Event plots, comparison between RNA load and probe load.** Experiments with Small RNA from H6914 (1st lot) or experiments with a probe. Consecutive 45 min long experiments with flow cell FAT05212 at -180mV (Table S2, Supplemental). Samples are 1 μL (blue trace), 2 μL (red trace), 3 μL (green trace), and 4 μL (yellow trace) of small RNA isolated from H6914, 1st lot, in ONT buffer and exhibit with no apparent concentration dependence. For comparison, an experiment is included with probe for miR-16 (rT4probe16, in Table 1) in buffer (blue dotted line) at 60,000 copies. Small RNA exhibits events with $(I_r/I_o)_{max}$ at 0.25 and at 0.35, but the event number due to small RNA is 4-fold less at $I_r/I_o \cong 0.25$, compared to the probe. A portion of the events at $I_r/I_o = 0.35$ is attributed to the ONT component(s) discussed in the text. The concentrations of the species in this experiment are comparable to the concentration of the species in all our experiments.

**Figure S3b: Event plots to illustrate detection or silencing of a probe for miR-16 in small RNA from H6914 (1st lot).** Consecutive experiments on flow cell FAS09254, each sample run for 45 min at -180mV (see Table S1 in the Supplementary). Events from the 1st and the 2nd runs were added to mimic a 90 min experiment. All three samples contain 44,100 copies of probe16AT to target miR-16 and variable amounts of small RNA isolated from H6914, at 0.1 μL (blue trace), 0.3 μL (red trace) and 0.2 μL (green trace). The small load of RNA was necessary in these experiments because miR-16 is highly expressed compared to all the other miRNAs we tested. Noticeably the blue and green traces include substantially more events at $(I_r/I_o)_{max}$ =0.20 compared to the red trace. In contrast to the late $I_r/I_o)_{max}$ =0.3 attributed to ONT component(s), the early $(I_r/I_o)_{max}$ is characteristic of the probe. Notably, the blue and green traces correspond to detection experiments, whereas the red trace corresponds to a silencing experiment.

The two experiments that “lie closer” to each other are the ones with 0.2 μL and 0.3 μL small RNA. The conclusion is drawn that miR-16 < 220,500 and miR-16 > 147,000, or due to the 2-fold concentration 73,500 < miR-16 < 110,250 in H6914. Additional experiments narrowed down this range to 100,000 < miR-16 < 110,250 (see Table S2 in Supplementary under flow cell FAT47395) using a different probe for miR-16.

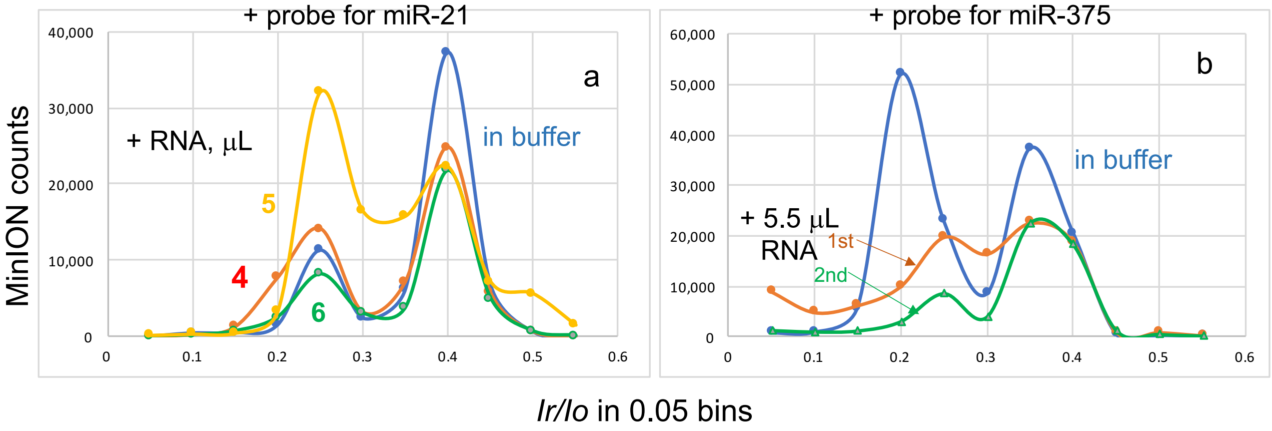

**Figure S4: Events plots to quantify (a) miR-21 and (b) miR-375 in small RNA isolated from H6914.**

**(a)** Four consecutive 45 min long experiments at -180mV using flow cell FAT14095 (Table S2 in Supplementary), with probe 21T5 (Table 1) at 57,240 copies. The first experiment contains only the probe 21T5 in buffer (blue trace (nonew)). The other three experiments include small RNA at 4 μL (red trace), at 6 μL (green trace) and at 5 μL (yellow trace). The last experiment clearly represents detection due to the high count of events at $(I_r/I_o)_{max}$ =0.25 and yields miR-21 < 11,448 from 57,240/5. Compared to the control (blue trace), the red trace represents a detection experiment because the Ratio late$(I_r/I_o)_{max}$ /early$(I_r/I_o)_{max}$ decreases, and yields miR-21 < 14,310 from 57,240/4. In turn, compared to the control (red trace), the green trace represents a silencing experiment with fewer events and an increase in Ratio late$(I_r/I_o)_{max}$ /early$(I_r/I_o)_{max}$ (see text and Protocol in Methods). The experiment represented by the green trace yields miR-21 > 9,540 from 57,240/6. These experiments are consistent and yield 9,540 < miR-21 < 11,448 or 14,310. The values closer to each other are 9,540 < miR-21 < 11,448 and arbitrarily post the average as copies of miR-21= 10,494 in small RNA or miR-21= 5,247 in 1μL serum H6914.
**(b)** Three 45 min long consecutive experiments were conducted with flow cell FAT89549 at -180mV (Table S2 in Supplementary). The first experiment illustrates the events plot of the probe m375T5 (Table 1) at 47,700 copies in buffer (blue trace (nonew)) and exhibits large peaks both at early $(I_r/I_o)_{max}$ =0.20 and at late $(I_r/I_o)_{max}$ =0.35. The source of these two peaks is discussed in the text. Second experiment with the same probe load and 5.5 μL small RNA isolated from H6914 1st lot exhibits substantially fewer events compare to the control with probe only, yielding a silencing experiment with miR-375 > 8,673 from 47,700/5.5 or miR-375 > 4,336 in 1μL serum H6914. Silencing is further confirmed by the 2nd run (green trace (nonew)), which exhibits even fewer events. Additional experiments with flow cell FAT05011 confirmed this result and yielded a range 4,200 < miR-375 <5,040 (Table S2 in Supplementary).

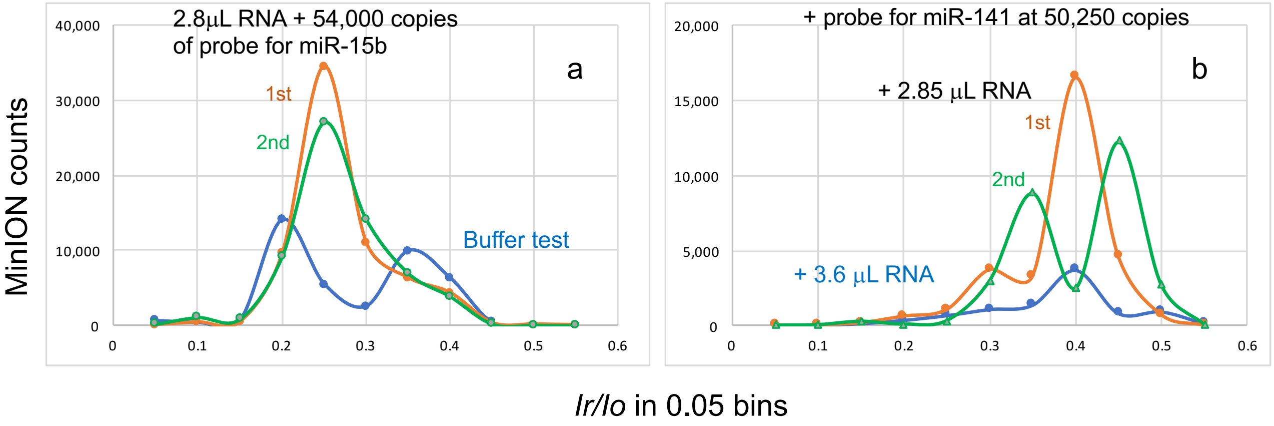

**Figure S5: Events plots from experiments to target miRNAs in small RNA isolated from the serum of a patient with (a) breast cancer or (b) prostate cancer (Table 2).**

**(a)** This figure represents a series of consecutive experiments done on flow cell FAU03318, all 45 min at -180mV. The control experiment here (blue trace, nonew) is a buffer test and exhibits two peaks at $(I_r/I_o)_{max}$ =0.2 and 0.35, a typical buffer profile. The next experiment is done with 2.8 μL from a Breast cancer small RNA sample and probe R15bT2 at a load of 54,000 copies (red trace). Total events in this experiment measure 1.5-fold higher compared to buffer; the 2$^{nd}$ run exhibits a similar profile (green trace (nonew)). There appears to be a shift from $(I_r/I_o)_{max}$ =0.2 to 0.25; shifts like that are not uncommon and are attributed to aging flow cells. Due to the higher number of events compared to the buffer (both in the 1$^{st}$ and in the 2$^{nd}$ run), this Breast cancer sample is conjectured to be a detection experiment with miR-15b < 9,634 from 54,000/2x2.8=9,634.

**(b)** This figure represents a series of experiments conducted for 45min at -180mV using flow cell FAT95653 carrying less than 200 active nanopores at this stage. The control experiment here is done with 3.6 μL from a Prostate cancer small RNA sample with probe 141T5 at a load of 50,250 copies (blue trace) reporting about 9,000 total events. Based on the earlier runs this experiment is considered a silencing experiment with miR-141 > 6,979 copies per 1 μL serum. The next experiment is done with a Prostate cancer sample with the same load of probe 141T5 but with a lesser small RNA load at 2.85 μL (red trace). Both the 1$^{st}$ and the 2$^{nd}$ run (green trace, nonew) with this sample exhibit each 3-fold more events compared to the control. Hence this sample suggests detection and yields miR-141 < 8,816 copies from 50,250/2x2.85=8,816 per 1 μL serum. The observations of dissimilar profiles, peak splitting, and peak shifting to a much later $(I_r/I_o)_{max}$ are attributed to an aging flow cell.

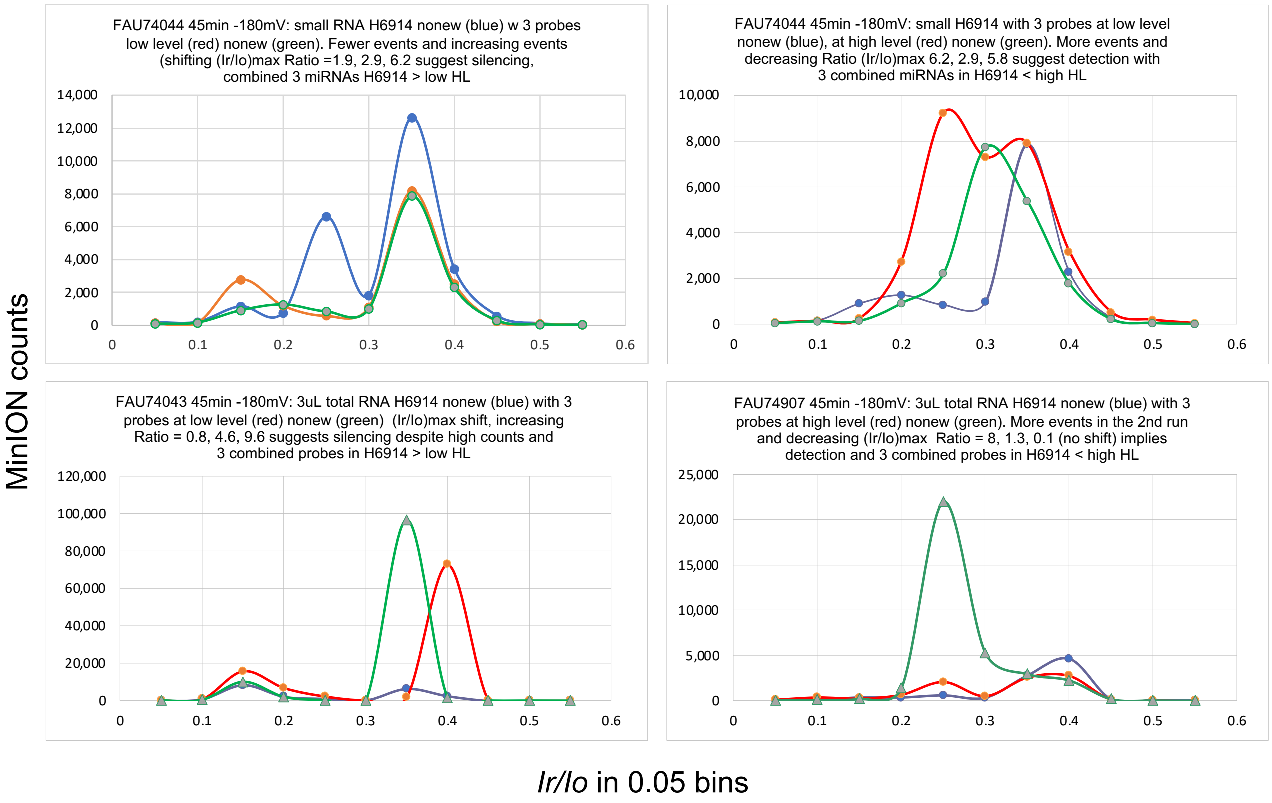

**Figure S6. Multiplexing 3 probes to target simultaneously miR-21, miR375 and miR-141 in the serum of H6914 2$^{nd}$ lot.** Samples were prepared with approximately 5uL of RNA, either small or total, and approximately 60,000 or 75,000 total probe copies used with small or total RNA, respectively. Probe count was matched to the miRNA level targeted for each different miRNA and accounted for the different concentration of each probe. Levels, LOW HL and HIGH HL, were the ones measured for H6914 1$^{st}$ lot (Table 2). In all figures blue trace is control, and red, green (nonew) are obtained from the test sample. Top left figure: experiment to target LOW HL from small RNA; silencing is observed compared to control. Top right figure: experiment to target HIGH HL from small RNA; detection is observed compared to control. Bottom left figure: experiment to target LOW HL from total RNA; they are more events compared to control, but these events exhibit a late (Ir/Io)max attributed to ONT components and the Ratio of late (Ir/Io)max/early (Ir/Io)max suggests silencing. Bottom right figure: experiment to target HIGH HL from total RNA; detection due to more events observed with the 2$^{nd}$ run and a decreased Ratio for both runs. These set of experiments the same quantitative conclusion from both total and small RNA obtained by the Monarch kit, namely the combined 3 probes for miR-21, miR-141 and miR-375 in H6914 (2$^{nd}$ lot) measure less than the HIGH HL and more than the LOW HL, as obtained from the sum of the individual miRNAs in H6914 (1$^{st}$ lot).

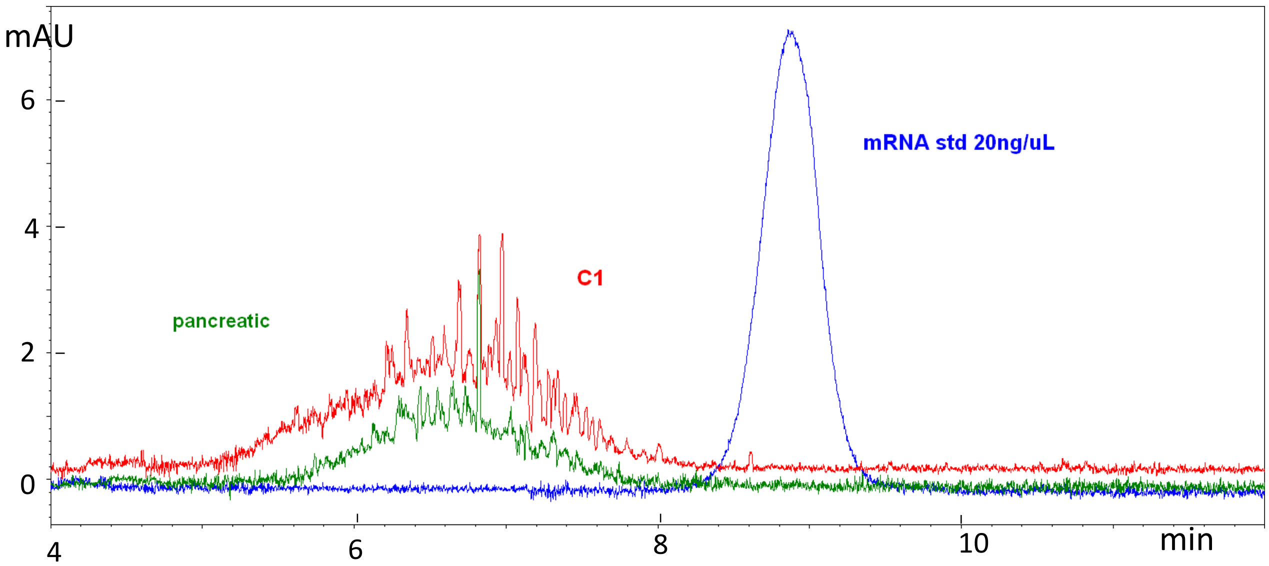

**Figure S7. Capillary Electrophoresis (CE) analyses at 260nm of total RNA obtained from serum using the Monarch RNA purification kit.** Concentration of samples in ng/μL is listed in Table 3 in text and was determined by comparing CE areas under the peak at 260nm between the unknown and the mRNA standard at 20ng/μL. It is noticeable that the standard is a sharper peak, while the RNA obtained from the serum samples is a much broader shoulder with many peaks, and this is because the standard is practically one component where the serum samples contain a plethora of RNA sequences composed. Due to the broadness of the peak LOQ is at 5 ng/μL (see Methods)

**Table S1: Serum samples from Discovery Life Sciences (DLS) with patient's logistics**

| ID,[1] Diagnosis, Cancer Stage | Disease Status | Treatment Status | Patient age at collection | Gender | Race |
|---|---|---|---|---|---|
| Pancreatic, II | Stable | Gemzar/Abraxane | 58 | Female | Black |
| Prostate, Adenocarcinoma, III | Newly Diagnosed | Pre-Treatment | 69 | Male | White |
| Breast, Invasive/Infiltrating Ductal, II | Newly Diagnosed | Pre-Treatment | 47 | Female | White |
| H1 | normal | NA | 36 | Female | Black |
| H2 | normal | NA | 67 | Male | White |
| H4 | normal | NA | 37 | Female | Black |
| Prostate CAN4 Adenocarcinoma, II | Newly Diagnosed | Pre-Treatment | 66 | Male | White |
| Prostate CAN6 Adenocarcinoma, II-b | Newly Diagnosed | Pre-Treatment | 56 | Male | White |
| Breast CAN9 Invasive/Infiltrating Ductal, I | Newly Diagnosed | Pre-Treatment | 58 | Female | White |

[1]First 3 entries from 1st DLS batch, the remainder from 2nd DLS batch. NA stands for not applicable.

### Table S2: Nanopore experiments conducted with the MinION, 45 min at -180mV

| Flow cell, sample | probe copies | total events | (Ir/Io)max | working channels /active pores | Result/Conclusion | miRNA copies | Fig # AVE/RSD |
|---|---|---|---|---|---|---|---|
| **FAS09254** | | | | | | | |
| buffe rtest | | 72,256 | 0.20 | | | | |
| nonew | | 31,415 | 0.30, 0.40 | | | | |
| 1uL smRNA H6914 | | 94,206 | 0.20 | | detection of 1uL small RNA at Ir/Io=0.20 | | |
| nonew | | 37,682 | 0.25 | | flow cell depleted of small RNA | | |
| nonew2 | | 49,434 | 0.30 | | " | | |
| 0.1uL smRNA + 5uL probe16AT 14.7fM (4.6 OsBp) | 44,100 | 49,385 | 0.30 | | events comparable to control/earlier experiment, | miR-16 < 220,500 | Fig. S3b |
| nonew | | 93,474 | 0.20, 0.30 | | double the events, many at Ir/Io=0.20, probe detection | | |
| nonew2 | | 59,684 | 0.20, 0.30 | | events at Ir/Io=0.2 cofirms detection | | |
| 0.3uL smRNA + 5uL probe16AT | 44,100 | 57,966 | 0.30 | | events cocparable to control and no events at Ir/Io=0.2, silencing | miR-16 > 73,500 | Fig. S3b |
| nonew | | 58,888 | 0.30 | 320/504 | silencing confirmed | | |
| 0.2uL smRNA + 5uL probe16AT | 44,100 | 70,940 | 0.20, 0.30 | 298/461 | more events compared to control and many at Ir/Io=0.2, detection | miR-16 < 110,250 | Fig. S3b |
| nonew | | 56,065 | 0.30 | 286/437 | unclear result | | |
| nonew2 | | 98,734 | 0.20, 0.30 | 272/404 | many events and some at Ir/Io=0.2, detection confirmed | 73,500 < miR-16 < 110,250 | |
| **FAT05212** | | | | | | | |
| buffer test | | 102,928 | 0.20, 0.35 | 469/1091 | | | |
| nonew | | 91,772 | 0.20, 0.35 | 466/1071 | | | |
| 1uL smRNA H6914 | | 66,597 | 0.35 | 462/1078 | many events observed with smRNA, including at Ir/Io=0.25 | | Fig. S3a |
| nonew | | 64,223 | 0.35 | 462/1067 | see Figure 3b, | | |
| 2uL smRNA | | 87,846 | 0.25, 0.35 | 461/1060 | some dependence of events on concentration if one adds both runs | | Fig. S3a |
| nonew | | 87,803 | 0.25, 0.35 | 462/1027 | | | |
| 3uL smRNA | | 102,298 | 0.20, 0.35 | 456/1033 | | | Fig. S3a |
| nonew | | 69,015 | 0.20 < 0.35 | 456/1046 | | | |
| 4uL smRNA | | 77,625 | 0.25 < 0.35 | 447/1005 | | | |
| nonew | | 96,695 | 0.25 << 0.35 | 442/996 | | | |
| 5uL rT4probe16 20fM in buffer | 60,000 | 82,130 | 0.25 = 0.35 | 418/943 | probe detection | | Fig. S3a |
| nonew | | 152,608 | 0.25>>0.35 | 433/957 | probe detection confirmed | | |
| nonew2 | | 88,077 | 0.25 = 0.35 | 443/961 | probe detection confirmed | | |
| 5uL small RNA | | 85,206 | 0.25<0.35 | 444/972 | | | |
| nonew | | 54,225 | 0.35 | 443/926 | | | |
| 5uL total RNA H6914 | | 157,423 | 0.25 | 395/797 | total RNA >> smRNA | | |
| nonew | | 126,497 | 0.20 > 0.25 | 405/747 | total RNA interfers with probe detection, many events at Ir/Io=0.20 | | |
| **FAT47395** | | | | | | | |
| buffer test, nonew2 | | 88,650 | 0.35 | 495/1244 | | | |
| 0.25uL smRNA + rT4probe16 | 60,000 | 181,568 | 0.35 | 494/1235 | more than 2x the events of the earlier run, detection | miR-16 < 120,000 | |
| nonew | | 133,967 | 0.05,0.2>0.35 | 453/1170 | detection confirmed | | |
| 0.30uL smRNA + rT4probe16 | 60,000 | 92,535 | 0.35>0.20, 0.05 | 489/1214 | reduced events and Ir/Io reversal, silencing | miR-16 > 100,000 | |
| nonew " | | 69,593 | 0.35>0.25 | 467/1190 | silencing confirmed | **100,000 < miR-16 < 110,250** | **105,125** |
| | | | | | | | 0.07 |
| clean, buffertest | | 53,426 | 0.35 | 480/1136 | | | |
| 7uL Rprobe15bT5 11.5 fM (1.5OsBp) in buffer | 48,300 | 71,345 | 0.2>0.25>0.35 | 408/957 | probe is detected, more counts and low Ir/Io | | |
| nonew " | | 59,173 | 0.25>0.2>0.35 | 464/1063 | buffer profile | | |
| 3ul smRNA + 3.5uL Rprobe15bT5 23fM | 48,300 | 60,743 | 0.2=0.3>0.35>0.25 | | same as buffer profile, silencing | miR-15b > 8,050 | |
| nonew | | 22,385 | 0.3>0.35>0.25 | | 3-fold fewer events, silencing confirmed | | |
| 2ul smRNA + 3.5uL Rprobe15bT5 | 48,300 | 47,373 | 0.25=0.2>0.3=0.35 | | 2-fold more events, detection | miR-15b < 12,075 | |
| nonew " | | 49,377 | 0.25>0.2=0.3>0.35 | | detection confirmed | | |
| 2.5uL smRNA + 3.5uL Rprobe15bT5 | 48,300 | 57,039 | 0.2>0.25 | 349/729 | same events as before, many events at low Ir/Io, detection | miR-15b < 9,660 | |
| nonew | | 66,036 | 0.3 | 330/683 | detection confirmed, two peaks came together | **8,050 < miR-15b < 9,660** | **8,855** |
| | | | | | | | 0.13 |
| **FAT43745** | | | | | | | |
| 2.5uL probe let7-b 36fM in buffer (6.1 OsBp) | 54,000 | 102,117 | 0.20>0.35 | 481/1174 | many events and at Ir/Io=0.2 clear detection of probe | | Fig. 2b |
| nonew | | 91,841 | 0.20>0.35 | 473/1136 | detection of probe | | Fig. 2b |
| 5uL smRNA + 2.5uL probe let7-b | 54,000 | 55,714 | 0.15,0.2<0.3,0.35 | 473/1127 | half of the events as compared to probe alone, silencing | let7-b > 5,400 | Fig. 2b |
| nonew " | | 49,850 | 0.15,0.2<0.3,0.35 | 461/1079 | silencing confirmed | | Fig. 2b |
| 4uL smRNA + 2.5uL probe let7b | 54,000 | 98,468 | 0.15,0.2>0.3 | 470/1085 | double the events compare detection at 4uL shRNA | let7-b < 6,750 | Fig. 2b |
| nonew " | | 44,368 | 0.30 | 461/1053 | | **5,400 < let7-b <6,750** | **6,075** |
| 2.5uL probe141T5 38.5fM in buffer, 6.9 | 57,750 | 81,945 | 0.20 | 437/989 | detection of probe | | 0.16 |
| nonew " | | 37,092 | 0.15=0.2=0.25=0.3 | 438/997 | low counts but at low Ir/Io consistent with probe | | |
| 10uL smRNA + 2.5uL probe141T5 | 57,750 | 44,078 | 0.2>0.3 | 434/966 | low counts, consistent with silencing | miR-141 > 2,888 | |
| nonew | | 42,973 | 0.2=0.25=0.3=0.35 | 427/944 | silencing confirmed | | |
| 7.5uL smRNA + 2.5uL probe141T5 | 57,750 | 122,792 | 0.25 | 429/968 | many events and at Ir/Io=0.25 clear detection of probe | miR-141 < 3,850 | |
| nonew | | 75,887 | 0.25 | 405/869 | events at low Ir/Io, detection confirmed | | |
| 9uL smRNA + 2.5uL probe141T5 | 57,750 | 94,249 | 0.1,0.2,0.25,0.35 | 421/865 | many events detection | miR-141 < 3,208 | |
| nonew | | 50,655 | 0.1,0.2,0.25,0.35 | 367/775 | events at low Ir/Io, detection confirmed | **2,888 < miR-141 < 3,208** | **3,048** |
| | | | | | | | 0.07 |
| **FAT05011** | | | | | | | |
| buffer test | | 60,943 | 0.25>0.2 | 473/1138 | 0.15 mL buffer 5-21-22 | | |
| nonew | | 34,327 | 0.25>0.2=0.35 | 471/1127 | | | |
| 3.5uL Rprobe15b 23fM (1.5 OsBp) in buffer | 48,300 | 41,884 | 0.25 | 464/1093 | number of events comparable to buffer, but Ir/Io=low | | |
| nonew | | 42,647 | 0.25 | 458/1066 | detection of probe confirmed but weak | | |
| 2.5uL smRNA + 3.5uL Rprobe15bT5 | 48,300 | 53,629 | 0.20=0.40 | 448/1023 | tentative detection due to Ir/Io=0.2 | miR-15b < 9,660 | |
| nonew | | 45,958 | 0.3 | 447/1008 | | | |
| 3.0uL smRNA + 3.5uL Rprobe15bT5 | 48,300 | 56,483 | 0.3 | 427/909 | tentative silencing due to Ir/Io=0.3 | miR-15b > 8,050 | |
| nonew | | 28,189 | 0.25=0.35 | 424/872 | | **8,050 < miR-15b < 9,660** | |
| 2.5uL probe375T5 33.6fM (5.0 OsBp) in buffe | 50,400 | 57,328 | 0.30>0.45 | 411/818 | probe detection, double the events compared to earlier run | | |
| 3uL smRNA + 2.5uL probe375T5 | 50,400 | 112,577 | 0.3>0.25>0.45 | 397/751 | double the events compared to earlier run, probe detection | miR-375 < 8,400 | |
| nonew | | 69,624 | 5 | 369/695 | | | |
| 6uL smRNA + 2.5uL probe375T5 | 50,400 | 42,778 | 0.35, 0.5 | 321/526 | only half the events, silencing | miR-375 > 4,200 | |
| 5uL smRNA + 2.5uL probe375T5 | 50,400 | 111,688 | 0.3>0.45 | 297/459 | double the events, detection | miR-375 < 5,040 | |
| 8uL smRNA + 2.5uL probe375T5 | 50,400 | 116,499 | 0.25=0.45 | 261/407 | double the events, detection | miR-375 > 3,150 | |
| | | | | | | **4,200 < miR-375 <5,040** | **4,620** |
| | | | | | | | 0.13 |
| **FAT12944** | | | | | | | |
| buffer test, nonew | | 25,007 | 0.25>0.35 | 411/814 | | | Fig. 2a |
| 3.5uL Rprobe15bT5 23fM (1.5 OsBp) in buff | 48,300 | 54,341 | 0.25>0.3>0.35 | 412/800 | double the events compared to buffer, probe detected | | Fig. 2a |

### Table S2, continues

| Flow cell, sample | probe copies | total events | (Ir/Io)max | working channels /active pores | Result/Conclusion | miRNA copies | Fig # AVE/RSD |
|---|---|---|---|---|---|---|---|
| **FAT14095** | | | | | | | |
| nonew buffer test | | 88,002 | 0.4 | 492/1263 | | | |
| 3uL probe21T5 31.8fM (6.2 OsBp) | 57,240 | 76,507 | 0.40>0.25 | 494/1281 | many events, probe detected even though Ir/Io high | | |
| nonew | | 67,690 | 0.40>0.25 | 493/1263 | detection confirmed by the many events | | Fig. S4a |
| 4uL smRNA + 3uL probe21T5 | 57,240 | 65,663 | 0.40>0.25>0.15 | 493/1262 | as many events as the probe in buffer, detection | miR-21 < 7,155 | Fig. S4a |
| 6uL smRNA + 3uL probe21T5 | 57,240 | 46,483 | 0.40>0.25 | 492/1220 | fewer events compared to earlier run, Ir/Io=0.4, silencing | miR-21 > 4,770 | Fig. S4a |
| 5uL smRNA + 3uL probe21T5 | 57,240 | 105,672 | 0.25>0.40 | 475/964 | double the events, Ir/Io=0.25, detection | miR-21 < 5,724 | Fig. S4a |
| | | | | | | **4,770 < miR-21 < 5,724** | **5,247** |
| | | | | | | | 0.13 |
| **FAT89549** | | | | | | | |
| buffer test | | 188,635 | 0.4 | 510/1579 | | | |
| nonew | | 148,404 | 0.4 | 509/1507 | | | |
| 3uL probe m375T5 26.5fM (4.4 OsBp) in buffe | 47,700 | 120,636 | 0.35<0.4 | 510/1502 | | | |
| " nonew | | 115,419 | 0.35<0.40 | 506/1486 | | | |
| 3uL probe m375T5 in buffer | 47,700 | 104,584 | 0.35>0.4=0.2 | 504/1473 | prepared new sample, detection of probe | | |
| nonew | | 148,568 | 0.2>0.35>0.25=0.4 | 500/1430 | confirmed | | Fig. S4b |
| 5.5uL smRNA(N/N) + 3uL probe m375T5 | 47,700 | 108,616 | 0.25=0.3,0.4 | 499/1414 | less events and reversal to high Ir/Io, silencing | miR-375 > 4,336 | Fig. S4b |
| nonew | | 61,261 | 0.35>0.4>0.25 | 487/1300 | silencing confirmed | | Fig. S4b |
| 4.5uL smRNA(N/N) + 3uL probe m375T5 | 47,700 | 61,713 | 0.4>0.35>0.25 | 485/1280 | unclear | | |
| nonew | | 102,462 | 0.25>0.3>0.35=0.4 | 477/1217 | many events, and reversal to low Ir/Io, detection | miR-375 < 5,300 | |
| | | | | | | **4,336 < miR-375 (N/N) < 5,300** | **4,818** |
| | | | | | | | 0.14 |
| 3.3uL PA smRNA + 3uL probe R15bT2 30fM (2.6 OsBp) | 54,000 | 68,651 | 0.2=0.25=0.4 | 427/899 | only 40% of the events of earlier exp, silencing | PA miR-15b > 8,182 | |
| nonew | | 45,720 | 0.20<0.40 | 391/763 | silencing confirmed | | |
| 2.8uL PA smRNA + 3uL probe R15bT2 | 54,000 | 71,142 | 0.3>0.4 | 355/680 | almost 2-fold the events of earlier run, detection | PA miR-15b < 9,643 | |
| nonew | | 76,479 | 0.3>0.35 | 333/600 | detection confirmed | **8,182 < PA miR-15b < 9,643** | **8,913** |
| buffer test | | 55,657 | 0.25>0.3 | 312/528 | | | **0.12** |
| 3.6uL BRE smRNA + 2.5uL probe141T5 33.5fM (6.9 OsBp) | 50,250 | 49,688 | 0.3 | 278/469 | events comparable to buffer test, silencing | BRE miR-141 > 6,979 | |
| nonew | | 41,197 | 0.35 | 263/434 | silencing confirmed but pores few | | |
| **FAT89764** | | | | | | | |
| 3uL RNA probe 16rT4 25fM (3.57 OsBp) | 45,000 | 58,422 | 0.35 | 354/893 | probe detectable at Ir/Io = 0.15 | | |
| nonew | | 55,969 | 0.35 | 353/874 | " | | Fig. S2 |
| **FAT94165** | | | | | | | |
| 3uL probe m21T5 in buffer, (4.4 OsBp) | 48,780 | 89,676 | 0.35>0.4 | 493/1337 | | | |
| nonew | | 107,917 | 0.25=0.35 | 491/1337 | probe detection confirmed | | |
| buffer test | | 67,979 | 0.35,0.4>0.2 | 496/1357 | good profile for buffer | | |
| 2.60uL PA smRNA + 3uL probe m375T5 26.5fM, 2x HL | 47,700 | 52,915 | 0.35>0.4 | 487/1316 | fewer events compared to buffer, silencing | PA miR-375 > 9,173 | |
| nonew | | 50,381 | 0.4>0.35 | 437/1056 | silencing confirmed | | Fig. 3a |
| 2.05uL PA smRNA + 3uL probe m375T5, 2.5x HL | 47,700 | 140,484 | 0.2, 0.4 | 410/945 | 3x the events compared to earlier run, detection | PA miR-375 < 11,634 | Fig. 3a |
| nonew | | 149,369 | 0.2>0.4 | 366/760 | detection confirmed | **9,173 < PA miR-375 < 11,634** | Fig. 3a, 3b |
| 4.75uL PA smRNA + 2.5uL probe141T5 33.5fM (4.7 OsBp), 1.7x HL | 50,250 | 25,093 | 0.35 | 351/701 | very few events, silencing | PA miR-141 > 5,289 | Fig. 3b |
| nonew | | 58,954 | 0.15, 0.3 | 277/509 | silencing confirmed, still 1/3 of the earlier sample | | Fig. 3b |
| 2.80uL PRO smRNA + 3uL probe miR15bT2 | 54,000 | 46,839 | 0.25, 0.30 | 281/496 | perhaps detection | PRO miR-15b < 9,643 | |
| nonew | | 69,396 | 0.2 | 268/457 | many events, confirmed detection | | |
| 3.30uL PRO smRNA + 3uL probe miR15bT2 | 54,000 | 48,858 | 0.15<0.40 | 260/431 | fewer events, silencing | PRO miR15b > 8,182 | |
| nonew | | 44,650 | 0.3 | 230/371 | confirmed silencing | | |
| | | | | | | **8,182 < PRO miR-15b < 9,643** | **8,913** |
| **FAT95653** | | | | | | | 0.12 |
| nonew buffer test | | 63,100 | 0.2>0.25=0.4 | 486/1315 | | | |
| 3.60uL PA smRNA + 2.5uL probe141T5, 2.3x | 50,250 | 75,181 | 0.35,0.4 | 484/1262 | tentative detection due to higher number of events | PA miR-141 < 6,979 | |
| nonew | | 111,048 | 0.25>0.30 | 344/651 | double the events compared to buffer, detection confirmed | **5,289 < PA miR-141 < 6,979** | |
| buffer test, nonew | | 86,617 | 0.2=0.25=0.55 | - | | | |
| 3.30uL BRE smRNA + 3uL probeR15bT2 | 54,000 | 40,176 | 0.25 | 277/507 | half ot the events compared to buffer, silencing | BRE miR-15b > 8,182 | |
| 2.85uL BRE smRNA + 2.5uL probe 141T5, 2.9x HL | 50,250 | 87,595 | 0.3 | 257/473 | double the events compared to earlier run, detection | BRE miR-141 < 8,816 | |
| nonew | | 68,809 | 0.25 | 236/435 | detection confirmed | **6,979 < BRE miR-141 < 8,816** | |
| 3.60uL PRO smRNA + 2.5uL probe 141T5, 2.3x HL | 50,250 | 15,593 | 0.3 | 226/381 | only 25% of the events of earlier run, silencing | PRO miR-141 > 6,979 | |
| nonew | | 9,416 | 0.4 | 197/333 | silencing confirmed | | Fig. S5b |
| 2.85uL PRO smRNA + 2.5uL probe 141T5, 2.9x HL | 50,250 | 30,859 | 0.4 | 191/295 | 3-fold more events compared to earlier run, detection | PRO miR-141 < 8,816 | Fig. S5b |
| nonew | | 30,081 | 0.45>0.35 | 174/271 | detection confirmed | | Fig. S5b |
| | | | | | | **6,979 < PRO miR-141 < 8,816** | |
| **FAU03300** | | | | | | | |
| probe R15bT2 in buffer | 54,000 | 40,053 | 0.35 | 494/1504 | unclear probe detection | | |
| nonew | | 46,521 | 0.35 | 492/1479 | " | | |
| 2.5uL probe 141T5 33.5fM (4.7 OsBp) in buffe | 50,250 | 106,992 | 0.3 | 488/1445 | detection but Ir/Io=0.3 for this flow cell | | |
| nonew | | 36,526 | 0.35 | 485/1412 | confirmed detection | | |
| 2.6uL BRE smRNA + 3uL probe m375T5 26.5fM (4.4 OsBp) | 47,700 | 44,603 | 0.15=0.2=0.35 | 482/1368 | detection based on low Ir/Io and somewhat higher events | BRE miR-375 < 9,173 | |
| nonew | | 87,949 | 0.35>0.3 | 467/1214 | confirmed detection | | |
| 3.45uL BRE smRNA + 3uL probe 375T5, 1.5xHL | 47,700 | 34,805 | 0.3 | 427/1032 | about 1/3 of the events from earlier fun, silencing | BRE miR-375 > 6,913 | |
| nonew | | 37,370 | 0.2>0.35 | 347/687 | similarly few events, silencing confirmed | **6,913 < BRE miR-375 < 9,173** | |
| 2.60uL PRO smRNA + 3uL probe 375T5 2xHL | 47,700 | 26,888 | 0.4>0.35>0.2 | 308/580 | to target prostate cancer miR375 at 9,173 | PRO miR-375 > 9,173 | |
| nonew | | 14,892 | 0.25=0.35 | 278/485 | silencing confirmed | | |
| 2.06uL PRO smRNA +3uL probe 375T5 2.5xHL | 47,700 | 108,032 | 0.25 | 241/414 | 7-fold the events observed in earlier run | PRO miR-375 < 11,578 | |
| nonew | | 63,650 | 0.25 | 214/369 | detection confirmed | **9,173 < PRO miR-375 < 11,578** | |

### Table S2, continues

| Flow cell, sample | probe copies | total events | (Ir/Io)max | working channels /active pores | Result/Conclusion | | | | | miRNA copies | Fig # AVE/RSD |
|---|---|---|---|---|---|---|---|---|---|---|---|
| **FAU03318** | | | | | | | | | | | |
| buffer test | | 120,449 | 0.4>0.2 | 492/1463 | | | | | | | |
| nonew | | 96,052 | 0.4>0.2 | 483/1449 | | | | | | | |
| 3uL probe R15bT2 30fM (2.6 OsBp) in buffer | 54,000 | 143,447 | 0.2>0.4 | 471/1418 | detection of probe due to more events and reversal of Ir/Io | | | | | | |
| nonew | | 106,414 | 0.2=0.4 | 470/1398 | detection of probe, confirmed | | | | | | |
| 3.5uL smRNA (N/N) +3uL probe R15bT2 | 54,000 | 66,118 | 0.4>0.35=0.2 | 460/1369 | half of the events compared to earlier run and Ir/Io reversal, silencing | | | | | miR-15b > 7,715 | |
| nonew | | 67,725 | 0.40, 0.15 | 452/1333 | silencing confirmed | | | | | | |
| 3.0uL smRNA (N/N) + 3uL probe R15bT2 | 54,000 | 69,990 | 0.2 | 447/1272 | detection due to Ir/Io=0.2 even though events same | | | | | miR-15b < 9,000 | |
| nonew | | 77,199 | 0.2 | 444/1190 | detection confirmed | | | | | **7,715 < miR-15b (N/N) < 9,000** | **8,358** |
| buffer test | | 39,935 | 0.2>0.25=0.35 | 427/1082 | | | | | | | 0.11 |
| nonew | | 40,651 | 0.2>0.35>0.4 | 417/1040 | | | | | | | Fig. S5a |
| 2.8uL BRE smRNA + 3uL probe R15bT2 | 54,000 | 66,812 | 0.25 | 409/986 | 50% more events compared to buffer, detection | | | | | BRE miR-15b < 9,643 | Fig. S5a |
| nonew | | 63,784 | 0.25 | 404/929 | detection, confirmed | | | | | **8,182 < BRE miR-15b < 9,643** | Fig. S5a |
| **FAU69853** | | | | | | | | | | | |
| buffer test | | 79,212 | 0.4>0.35 | 507/1535 | | | | | | | |
| nonew | | 84,991 | 0.4>0.35, 0.25 | 504/1525 | | | | | | | |
| 3.1uL C1 smRNA +3uL pr m21T5 27.1fM | 48,780 | 54,303 | 0.4 | 499/1480 | probe to target 1.5x HL | | | | | | |
| nonew | | 50,232 | 0.4 | 440/1116 | silencing | | | | | C1 miR-21 > 7,868 | |
| 2.3uL C1 smRNA +3uL pr m21T5 27.1fM | 48,780 | 25,589 | 0.4 | 409/965 | probe to target 2x HL | | | | | | |
| nonew | | 28,305 | 0.4 | 328/731 | silencing confirmed | | | | | C1 miR-21 > 10,604 | |
| buffer test | | 22,270 | 0.25,0.35,0.4 | 301/629 | | | | | | **C1 miR-21 > 2x HL** | Table 2 |
| buffer test | | 19,976 | 0.35>0.4 | 354/664 | | | | | | | |
| 3uL total RNA CAN4 | | 15,996 | 0.35>0.4 | 360/686 | patient ID: prostate CAN4 | | | | | | |
| nonew | | 15,718 | 0.35>0.4 | 337/629 | | | | | | | |
| 3uL total RNA CAN4 + 2x HL (3 probes combined) | | 20,628 | 0.2>0.35 | 349/644 | detection in the 1st run | | | | | CAN4 (prostate) < 2x HL (3 probes) | |
| nonew | | 10,088 | 0.35>0.4 | 327/585 | | | | | | | |
| 3uL totRNA breast cancer + 2x HL (3probes) | | 23,328 | 0.35>0.4 | 299/520 | detection in the 1st run | | | | | breast cancer < 2x HL (3 probes) | |
| nonew | | 6,489 | 0.35>0.4 | 276/454 | | | | | | | |
| buffer test | | 17,093 | 0.4>0.35 | 257/412 | | | | | | | |
| 5.6uL total RNA prostate CAN 6 | | 27,320 | 0.4 | 397/736 | | | sum of 1st and 2nd runs | | | | |
| nonew | | 9,431 | 0.2 | 262/458 | | | 36,751 | | | | Fig. 4 |
| 5.6uL total RNA CAN 6 + 4uL probe15bT5 | 42,000 | 10,479 | 0.45>0.2 | 199/313 | silencing | | | | | | |
| nonew | | 10,515 | 0.4>0.2 | 96/126 | silencing confirmed | | 20,994 | | | prostate CAN6 miR15b >7,500 | Fig. 4 |
| 5uL total RNA CAN 6 + 4uL probe15bT5 | 42,000 | 27,412 | 0.2 | 85/93 | detection with 20 pores | | | | | | |
| nonew | | 47,018 | 0.2>0.4 | 48/52 | detection confirmed | | 74,430 | | | prostate CAN6 miR15b <8,400 | Fig. 4 |
| | | | | | | | | | | **7,500 < prostate CAN 6 <8,400** | **7,950** |
| | | | | | | | | | | | 0.08 |
| **FAU74044** | | | | | | | | | | | |
| buffer test | | 74,112 | 0.4 | 510/1560 | | | | | | | |
| nonew | | 77,278 | 0.4 | 508/1547 | | | | | | | |
| 3uL probe let7bT5 30fM (6.1 OsBp) | 54,000 | 177,226 | 0.25>0.4 | 509/1529 | probe detection in 1st run | | | | | | |
| nonew | | 86,915 | 0.4 | 506/1508 | | | | | | | |
| 2.2uL BRE smRNA + 3uL pro let7bT5 2x HL | 54,000 | 85,107 | 0.4>0.35 | 508/1486 | detection, events as many as probe alone | | | | | | |
| nonew | | 89,834 | 0.35>0.4 | 489/1317 | detection confirmed | | | | | **BRE let7-b < 9,121, 12,272** | |
| 2.96uL BREsmRNA + 3uL pro let7bT5 1.5xHl | 54,000 | 170,113 | 0.35=0.15 | 458/1178 | detection, more events, (Ir/Io)max reversal | | | | | **BRE let7-b < 1.5 x HL, 2.0 x HL** | |
| nonew | | 81,028 | 0.15>0.4 | 418/1034 | | | | | | | |
| 2.32uL PAN smRNA + 3uL pro miR21 2x HL | 57,240 | 46,915 | 0.35>0.4>0.25=0.2 | 399/945 | silencing, half of the events of the control | | | | | **PAN miR-21 > 12,336** | |
| nonew | | 45,873 | 0.35>0.4, 0.2 | 360/844 | silencing confirmed | | | | | **PAN miR-21 > 2.0 x HL** | |
| buffer test | | 39,049 | 0.35>0.25 | 393/895 | | | | | | | |
| 3uL small RNA H6914 2nd lot | | 37,881 | 0.25=0.35 | 387/835 | R 0.35/0.25=1.12 | | | | | | |
| nonew | | 27,508 | 0.25<0.35 | 363/788 | R 0.35/0.25=1.91 | | | | | | |
| 3probes LOW HL + small RNA H6914 2nd lo | 36,605 | 17,056 | 0.35>0.15 | 355/769 | R 0.35/0.15=2.94 | | silencing | | | | Fig. S6 |
| nonew | | 14,836 | 0.35 | 333/721 | R 0.35/0.2=6.2 | | silencing confirmed | | | | Fig. S6 |
| 3 probes HIGH HL + small RNA H6914 2nd l | 43,926 | 31,589 | 0.25>0.35>0.3 | 332/706 | (Ir/Io)max reversal | | detection | | | | Fig. S6 |
| nonew | | 18,716 | 0.3 | 307/670 | " | | detection confirmed | | | | Fig. S6 |
| | | | | **3 probes for 3 targets = miR-21 + miR-375 + miR-141** | | | | | | **LOW HL < 3 probes small RNA H6914 < HIGH HL** | |
| buffer test | | 41,920 | 0.4 | 363/735 | | | | | | | |
| 4.8uL totRNA H6914 3rd lot | | 59,699 | 0.35,04,03,015 | 418/907 | | 80,040 | | | | | |
| nonew | | 20,341 | 0.45 | 360/765 | | | | | | | |
| 4.8uL totRNA H4522 + 4uL probe15bT5 17.5fM | 42,000 | 56,449 | 0.3 | 343/685 | detection, more events | | | | | | |
| nonew | | 35,516 | 0.4>0.45>0.25 | 319/631 | detection confirmed | | | | | **total RNA H4522, miR-15b < 8,750** | |
| 5.2uL totRNA H4522 + 4uL probe15bT5 | 42,000 | 126,114 | 0.35 | 290/565 | detection , more events | | | | | **total RNA H4522, miR-15b < 8,077** | Fig. S6 |
| nonew | | 106,818 | 0.35 | 257/474 | detection confirmed | | | | | | |
| 6uL total RNA H4522 + 4uL probe15bT5 | 42,000 | 16,370 | 0.3>0.25 | 256/442 | silencing | | | | | **total RNA H4522, miR-15b > 7,000** | Fig. S6 |
| nonew | | 68,768 | 0.45>0.25 | 241/400 | silencing confirmed | | | | | **7,000< total RNA H4522 miR-15b < 8,077** | |
| | | | | | | | | | | | **total RNA H4522, miR-15b = 7,539** |
| | | | | | | | | | | | 0.10 |
| 4.8uL totRNA CAN4 | | 30,343 | 0.3 | 230/362 | control | | | | | | |
| nonew | | 22,581 | 0.3 | 195/302 | comparable counts in all experiments | | | | | | |
| 4.8uL totRNA CAN4 + 5uL probe15bT5 | 52,500 | 42,626 | 0.3, 0.4 | 152/216 | silencing, material shifts to later Ir/Io | | | | | **total RNA CAN4 miR-15b > 10,938** | |
| nonew | | 26,226 | 0.35 | 162/216 | confirmed silencing | | | | | | |
| 4.2uL totRNA CAN4 + 5uL probe15bT5 | 52,500 | 22,270 | 0.35 | 186/240 | detection, material shifts to earlier Ir/Io | | | | | **total RNA CAN4 miR-15b < 12,500** | |
| nonew | | 31,949 | 0.3 | 136/170 | | | | | | | **total RNA CAN4 miR-15b = 11,719** |
| | | | | | | | | | | | 0.09 |
| **FAU64633** | | | | | | | | | | | |
| buffer test | | 86,610 | 0.35>>0.4 | 507/1625 | | | | | | | |
| nonew | | 81,390 | 0.35>0.4 | 507/1626 | | | | | | | |
| 2.84uL small RNA C1+ 3uL probe R15bT2 30fM 2.6Os | 54,000 | 60,377 | 0.35>0.4 | 507/1594 | fewer counts, silencing | | | | | C1 miR-15b > 9,507 | |
| nonew | | 76,018 | 0.15,0.2,0.35,0.4 | 418/974 | rejected due to the high loss of pores , | | | | | | |
| 2.50uL small RNA C1 + 3uL probe R15bT2 | 54,000 | 54,818 | 0.4>0.35, 0.25 | 344/711 | fewer counts, silencing | | | | | | |
| nonew | | 29,686 | 0.35>0.4 | 260/496 | silencing confirmed | | | | | **C1 miR-15b > 10,800** | Table 2 & 3 |
| buffertest | | 31,527 | 0.35>0.3 | 192/346 | | | | | | | |

### Table S2, continues

| Flow cell, sample | probe copies | total events | (Ir/Io)max | working channels /active pores | Result/Conclusion | | miRNA copies | Fig # AVE/RSD |
|---|---|---|---|---|---|---|---|---|
| **FAU74979** | | | | | | | | |
| buffer test | | 61,247 | 0.4 | 499/1545 | | | | |
| nonew | | 73,966 | 0.4>>0.2 | 500/1526 | | | | |
| 3uL probe let7bT5 (30fM, 6.1 OsBp) | 54,000 | 83,244 | 0.4 | 496/1499 | probe detection by more events | | | |
| nonew | | 86,735 | 0.4 | 497/1483 | | | | |
| pro let7bT5 + 2.20uL prostate smRNA 2x HL | 54,000 | 70,704 | 0.4>>0.35 | 486/1433 | detection, comparable to probe | | **prostate small RNA let7-b < 2x HL** | |
| nonew | | 73,782 | 0.4>>0.35 | 479/1346 | " | | | |
| pro let7bT5 + 2.96uL prostate smRNA 1.5xHL | 54,000 | 51,581 | 0.4>0.3,0.35 | 459/1199 | silencing, fewer events | | **prostate small RNA > 1.5 x HL** | |
| nonew | | 46,330 | 0.4>0.3 | 386/874 | silencing confirmed | | **1.5x HL < prostate small RNA let7-b < 2 x HL** | |
| pro let7bT5 + 2.22uL pancreatic smRNA 2x HL | 54,000 | 60,677 | 0.4=0.3, 0.25, 0.2 | 338/625 | detection | | | |
| nonew | | 134,616 | 0.3 | 249/427 | detection confirmed | | **pancreatic small RNA let7-b < 2x HL** | |
| buffer test | | 146,945 | 0.3>0.35>0.4 | 294/481 | | | | |
| 3uL small RNA H6914 2nd lot | | 129,039 | 0.3>0.35 | 262/421 | Ratio late/early (Ir/Io)max (0.35/0.3)=0.68 | | | Fig. S6 |
| nonew | | 93,058 | 0.35>0.3,0.4 | 243/379 | R 0.35/0.3=1.7 | control run | | |
| 3probes LOW HL + 3uL small RNA H6914 | 36,605 | 49,618 | 0.35, 0.3 | 245/373 | R 0.35/0.3=8.2 | higher R means silencing | | |
| nonew | | 27,180 | 0.35 | 230/344 | R 0.35/0.3=12.7 | confirmed, fewer counts too/control run for the next | | |
| 3probes HIGH HL + 3uL small RNA H6914 | 43,926 | 8,540 | 0.4>0.35>0.3 | 227/335 | R 0.35/0.3=1.4 | lower R, implies detection | **LOW HL < small RNA H6914 (2nd lot) 3 probes < HIGH HL** | |
| nonew | | 26,515 | 0.35 | 212/313 | R 0.35/0.3=3 | confirmed, higher counts in 2nd run | **repeat exp, see FAU74044** | |
| buffer test | | 78,217 | 0.35 | 226/329 | | | | |
| 6.2uL totRNA CAN 9 | | 28,594 | 0.35 | 297/469 | control totRNA | | | |
| nonew | | 14,653 | 0.25>0.4>0.3=0.35 | 177/254 | | | | |
| 6.2uL totRNA CAN 9 + 5uL probe15bT5 | 52,500 | 18,996 | 0.45 | 162/216 | silencing | | | |
| nonew | | 2,343 | 0.4 | 72/86 | silencing confirmed, shift to higher Ir/Io | | **total RNA CAN 9 miR-15b > 8,468** | |
| 5.6uL totRNA CAN 9 + 5uL probe15bT5 | 52,500 | 1,296 | 0.35 | 52/61 | detection, shift to lower Ir/Io | | | |
| nonew | | 1,105 | 0.35 | 29/30 | " | | **total RNA CAN 9 miR-15b < 9,375** | |
| | | | | | | | **total RNA CAN 9 miR-15b =** | **8,922** |
| | | | | | | | | 0.07 |
| **FAU74907** | | | | | | | | |
| buffer test | | *66,509* | 0.4>0.35 | 509/1677 | | | | |
| nonew | | 71,041 | 0.4>0.35 | 510/1671 | | | | |
| 3.4uL smRNA C1 +3uL pro m375T5 1.5x HL | 47,700 | 47,588 | 0.4 | 509/1654 | silencing, fewer probes | | **small RNA C1 miR-375 > 1.5x HL** | |
| nonew | | 43,921 | 0.4 | 387/823 | | | | |
| 2.6uL smRNA C1 +3uL pro m375T5 2x HL | 47,700 | 27,370 | 0.4, 0.15 | 297/596 | detection, reversal early (Ir/Io)max | | **small RNA C1 miR-375 < 2x HL** | |
| nonew | | 18,712 | 0.4 | 221/402 | | | **1.5x HL < small RNA C1 miR-375 < 2x HL** | |
| buffer test | | 27,971 | 0.3>0.4 | 212/352 | | | | |
| buffer test | | 13,966 | 0.4>0.35 | 291/450 | | | | |
| 3uL total RNA H6914 2nd lot | | 13,107 | 0.4>0.35>0.25 | 308/493 | | | | |
| nonew | | 9,422 | 0.4>0.35 | 251/392 | | | | Fig. S6 |
| 3 probes HIGH HL w tot H6914 2nd lot | 43,926 | 9,684 | 0.25=0.35=0.4 | 239/372 | | | | Fig. S6 |
| nonew | | 34,465 | 0.25 | 211/327 | detection, more events & material shift to early (Ir/Io)max | | **total RNA H6914 3 probes < HIGH HL** | Fig. S6 |
| 3uL total RNA C1 | | 8,826 | 0.35>0.4,0.25 | 217/339 | | | | |
| nonew | | 13,067 | 0.3>0.35 | 178/281 | | | | |
| 3probes 2x HL w 3uL total RNA C1 (repeat) | | 36,659 | 0.35 | 199/307 | detection, more events | | | |
| nonew | | 47,082 | 0.35 | 174/255 | detection confirmed | | **total RNA C1 3 probes < 2.0x HL** | |
| buffer test | | 2,206 | 0.35>0.4 | 161/217 | | | | |
| 8uL total RNA H4522 | | 49,256 | 0.40 | 408/757 | | | | |
| nonew | | 40,596 | 0.40 | 183/260 | | | | |
| 8uL total RNA H4522+ 2uL probe m21T5 27.1fM | 32,520 | 23,331 | 0.40 | 155/188 | silencing, fewer probes | | **total RNA H4522 lot miR-21 < 4,065** | |
| nonew | | 13,556 | 0.35 | 108/126 | " | | | |
| **FAU74043** | | | | | | | | |
| buffer test | | 92,397 | 0.4=0.35 | 512/1668 | after adding 0.6mL ONT buffer filtered 7-10 and removing excess | | | |
| nonew | | 102,104 | 0.35>0.4>>0.2 | 511/1679 | Ratio late/early (Ir/Io)max: R=2 | | | |
| 2.2uL probe 141T5 (38.5fM) +5.6uL smRNA C1 1.5x HL | 50,820 | 70,866 | 0.4 & low Ir/Io | 511/1671 | R=4.2 silencing, fewer events and increasing Ratio | | | |
| nonew | | 55,180 | 0.35=0.4 | 337/744 | R=12.3 silencing, fewer events and increasing Ratio | | | |
| 2.2uL probe 141T5 +4.2uL smRNA C1 2x HL | 50,820 | 47,130 | 0.35>0.4, 0.2 | 234/457 | R=6.9 detection, comparable events but decreasing Ratio | | **1.5x HL < small RNA C1 miR-141 < 2.0 HL** | |
| nonew | | 54,554 | 0.25 | 146/255 | R=0.2 detection, comparable events but decreasing Ratio | | | |
| buffer test | | 5,932 | 0.4>0.35 | 117/175 | | | | |
| buffer test | | 27,567 | 0.4, 0.2 | 253/350 | | | | |
| 3uL total RNA H6914 2nd lot | | 34,995 | 0.2=0.35, 0.15>0.4 | 245/350 | | | | |
| nonew | | 22,177 | 0.15>0.35 | 148/208 | | | | Fig. S6 |
| 3probes LOW HL + 3uL total RNA H6914 | 36,605 | 101,236 | 0.4>>0.15 | 142/188 | silencing, material shift to late Ir/Io | | | Fig. S6 |
| nonew | | 111,599 | 0.35 | 123/157 | " | | **LOW HL < total RNA H6914 (2nd lot) 3 probes** | Fig. S6 |
| **FAU74891** | | | | | | | | |
| buffer test | | 58,764 | 0.4 | 511/1678 | | | | |
| nonew | | 61,869 | 0.4>0.35>0.3 | 511/1674 | | | | |
| 3uL probe m21T5 27.1fM | 48,780 | 72,527 | 0.4>0.35 | 510/1662 | | | | |
| nonew | | 71,894 | 0.4>0.35 | 511/1644 | | | | |
| 3uL probe m21T5 + 2.3uL PRO smRNA 2xHL | 48,780 | 73,602 | 0.4 | 511/1633 | detection, events like w probe alone | | PRO small RNA miR-21 < 2x HL | |
| nonew | | 66,729 | 0.4>0.35>0.3 | 508/1614 | " | | | |
| 3uL pro m21T5 + 3.1uL PRO smRNA 1.5xHL | 48,780 | 46,069 | 0.4>0.35 | 507/1599 | silencing, fewer events | | PRO small RNA miR-21 > 1.5x HL | |
| nonew | | 46,492 | 0.4>0.35 | 493/1402 | " | | **1.5x HL < PRO small RNA miR-21< 2x HL** | |
| 3uL pro m21T5 + 1.33uL BRE smRNA 3.5xHl | 48,780 | 57,399 | 0.4>0.35, 0.15 | 464/1228 | detection | | | |
| nonew | | 43,582 | 0.4>0.35 | 437/1118 | | | | |
| 3uL pro m21T5 + 1.16uL BRE smRNA 4xHL | 48,780 | 49,179 | 0.25=0.35=0.4 | 422/1034 | detection | | **BRE small RNA miR-21 < 3.5, 4x HL** | |
| nonew | | 27,303 | 0.35>0.4, 0.25 | 380/920 | | | | |

### Table S2, continues

| Flow cell, sample | probe copies | total events | (Ir/Io)max | working channels /active pores | Result/Conclusion | | | | | miRNA copies | Fig # AVE/RSD |
|---|---|---|---|---|---|---|---|---|---|---|---|
| **FAU70015** | | | | | | | | | | | |
| buffertest | | 90,759 | 0.4 | 506/1579 | | | | | | | |
| nonew | | 92,818 | 0.4 | 503/1564 | | | | | | | |
| 3uL probe m21T5 27.1fM 4.4 OsBp | 48,780 | 127,628 | 0.4>0.45 | 503/1552 | probe detection, more events | | | | | | |
| nonew | | 78,651 | 0.4>>0.35 | 503/1539 | control | | | | | | |
| 3uL pro m21T5 + 2.3uL BRE smRNA 2xHL | 48,780 | 59,213 | 0.4 | 500/1521 | silencing, fewer events | | | | | **BRE miR-21 > 2x HL** | |
| nonew | | 64,746 | 0.4 | 492/1437 | control; Ratio=16 | | | | | | |
| 3uL pro m21T5 + 1.86uL BRE smRNA 2.5xHl | 48,780 | 47,900 | 0.4 | 488/1365 | detection Ratio decreases R=13 | | | | | **BRE miR-21 < 2.5, 3.0x HL** | |
| nonew | | 52,934 | 0.4>0.25 | 472/1195 | detection confirmed, R=3 | | | | | | |
| 3uL pro m21T5 + 1.55uL BRE smRNA 3xHL | 48,780 | 44,428 | 0.4>0.35=0.25 | 450/1053 | detection, material shift | | | | | **2x HL <BRE small RNA miR-21 < 2.5x HL** | |
| nonew | | *84,703* | 0.2,0.25,0.4,0.5 | 423/918 | detection confirmed | | | | | | |
| buffertest | | 58,808 | 0.3 | 332/573 | | | | | | | |
| 3uL PRO total RNA | | 28,947 | 0.3>0.35>0.4 | 360/607 | | | | | | | |
| nonew | | 11,871 | 0.4>0.35>0.3 | 309/513 | | | | | | | |
| 3probes 2x HL w 3uL totRNA PRO | | 10,094 | 0.2=0.25=0.35=0.4 | 306/496 | detection, material shift to early Ir/Io | | | | | **3 probes total RNA PRO < 2x HL** | |
| nonew | | 13,722 | 0.45 | 269/424 | | | | | | | |
| buffertest | | 16,439 | 0.25 | 272/408 | | | | | | | |
| 5.6uL total RNA CAN 6 | | 40,483 | 0.4 | 379/656 | | duplicate w FAU69853 | | | | | |
| nonew | | 9,425 | 0.3>0.4 | 276/460 | | | | | | | |
| 5.6uL total RNA CAN 6 + 4uL probe15bT5 | 42,000 | 21,120 | 0.25>0.35>0.4 | 252/398 | silencing , material shift to early Ir/Io | | | | | | |
| nonew | | 26,788 | 0.4 | 229/358 | " | | | | | 7,500 < tot RNA CAN 6 miR-15b < 8,400 | |
| 5uL total RNA CAN 6 + 4uL probe15bT5 | 42,000 | 58,017 | 0.35 | 217/319 | detection more events | | | | | **total RNA CAN 6 miR-15b =** | **7,950** |
| nonew | | 71,301 | 0.35=0.4 | 193/266 | detection confirmed miR-15b < 8,400 | | | | | | 0.08 |
| **FAV43661 (R10)** | | | | | | | | | | | |
| 4.0uL totRNA CAN4 | | 103,141 | 0.15>0.10=0.3>0.35 | 269/393 | | | | | | | |
| nonew | | 113,666 | 0.15>0.10=0.3>0.35 | 236/335 | control | | | | | | |
| 4.0uL totRNA CAN4 + 2uL probe15bT5 | 21,000 | 74,662 | 0.15 | 113/132 | silencing, fewer counts | | | | | **total RNA CAN 4 miR-15b > 5,250** | |
| nonew | | 69,664 | 0.15 | 202/253 | silencing confirmed | | | | | | |
| 4.8uL smallRNA H4522 | | 98,118 | 0.05,0.15 | 184/226 | | | | | | | |
| nonew | | 49,323 | 0.05 | 140/170 | control | | | | | | |
| 4.8uL smallRNA H4522 + 4uL probe15bT5 | 42,000 | 46,161 | 0.05 | 63/69 | detection, few events due to very few pores | | | | | miR-15b smRNA H4522 < 8750 | |
| nonew | | 34,592 | 0.05 | 88/102 | control | | | | | | |
| 5.2uL smallRNA H4522 + 4uL probe15bT5 | 42,000 | 65,099 | 0.2>0.05>0.25 | 24/26 | detection ( Ir/Io)max at 0.2 | | | | | miR-15b smRNA H4522 < 8077 | |
| nonew | | 64,465 | 0.2>0.05 | 46/53 | control | | | | | | |
| 6uL smallRNA H4522 + 4uL probe15bT5 | 42,000 | 81,411 | 0.05=0.2=0.25 | 79/86 | comparable to control | | | | | miR-15b smRNA H4522 > 7000 | |
| nonew | | 19,242 | 0.2 | 96/105 | silencing confirmed | | | | | **H4522 small RNA miR-15b =** | **7,539** |
| | | | | | | | | | | | 0.10 |
| **FAV43399 (R10)** | | | | | | | | | | | |
| buffertest | | 137,626 | 0.2>0.15>0.35 | 238/359 | | | | | | | |
| nonew | | 107,403 | 0.35=0.2>0.15 | 239/344 | | | sum of 1st and 2nd | | | | |
| 7.7uL small RNA H2 | | 113,069 | 0.15>0.2>0.35 | 220/321 | | | | | | | |
| nonew | | 93,840 | 0.15=0.2>0.35 | 183/273 | control | | 206,909 | | | | |
| 7.7uL small RNA H2 + 4uL m21T5 27.1fM | 65040 | 79,533 | 0.35>0.10 | 179/256 | unclear | | | | | | |
| nonew | | 55,362 | 0.15>0.35 | 164/229 | control for next | | 134,895 | | | | |
| 8.2uL small RNA H2 + 4uL m21T5 | 65040 | 52,610 | 0.2 | 137/175 | | | | | | | |
| nonew | | 154,358 | 0.2 | 101/131 | detection | | 206,968 | | | **small RNA H2 miR-21 < 7,932** | |
| **FAV45860 (R10)** | | | | | | | | | | | |
| buffertest | | 235,653 | 0.15=0.2 | 385/830 | | | | | | | |
| nonew | | 163,038 | 0.15>0.3>0.2 | 370/773 | control buffer | | Sum of 1st + 2nd run | | | | |
| 6.2uL small RNA H1 | | 172,364 | 0.15>0.2>0.1 | 357/720 | control RNA | | | | | | |
| nonew | | 113,475 | 0.15>0.3>0.35 | 319/599 | | | 285,839 | control | | | |
| 6.2uL small RNA H1 + 4uL m21T5 27.1fM | 65,040 | 131,271 | 0.15>0.2>0.35 | 278/506 | detection, material shift | | | | | small RNA H1 miR-21 < 10,490 | Fig. 5 |
| nonew | | 115,676 | 0.15 | 259/412 | to early Ir/Io | | 246,947 | detection | | | Fig. 5 |
| 7.2uL small RNA H1 + 4uL m21T5 27.1fM | 65,040 | 62,530 | 0.15>0.35 | 212/318 | silencing, few counts | | | | | small RNA H1 miR-21 > 9,033 | Fig. 5 |
| nonew | | 47,593 | 0.15>0.2=0.35 | 176/268 | " | | 110,123 | silencing | | **9,033 < H1 smRNA miR-21 < 10,490** | **9,762** |
| 4uL probe m21T5 new dilution | | 37,920 | 0.15>0.10>0.35 | 152/200 | | | | | | | 0.11 |
| nonew | | 34,290 | 0.15 | 81/98 | | | 72,210 | control | | | |
| 4uL probe m21T5 + 8uL tot RNA CAN4 | 65,040 | 74,098 | 0.15 | 189/232 | detection, more events | | | | | total RNA CAN 4 miR-21 < 8,130 | |
| nonew | | 71,955 | 0.15>0.2=0.4 | 255/354 | | | 146,053 | detection | | | |
| 4uL probe m21T5 + 9uL tot RNA CAN4 | 65,040 | 40,240 | 0.25 | 125/147 | silencing, fewer events and shift | | | | | total RNA CAN 4 miR-21 > 7,227 | |
| nonew | | 78,843 | 0.05=0.25 | 182/239 | | | 119,083 | silencing | | **7,227 < CAN4 totRNA miR-21 < 8,130** | **7,679** |
| | | | | | | | | | | | 0.08 |
| **FAS88208** | | | | | | | | | | | |
| buffer test | | 105,365 | 0.35>0.4 | 510/1671 | | | | | | | |
| nonew | | 105,674 | 0.35>0.4 | 510/1665 | | | | | | | |
| 3uL small RNA H1 | | 85,736 | 0.35>>0.4 | 501/1618 | | | | | | | |
| nonew | | 75,682 | 0.35 | 465/1252 | | | | | | | |
| 3uL smRNA H1 + 4uL probe 15bT5 17.5fM | 42,000 | 38,371 | 0.35 | 401/972 | silencing due to less events | | | | | small RNA H1 miR-15b > 14,000 | |
| nonew | | 30,610 | 0.35 | 300/583 | | | | | | | |
| 3uL smRNA H1 + 4.4uL probe 15bT5 | 46,200 | 28,272 | 0.3>0.35 | 226/388 | silencing due to less events | | | | | small RNA H1 miR-15b > 15,400 | |
| nonew | | 9,818 | 0.35 | 148/223 | R=24 | | | | | | |
| 3uL smRNA H1 + 4.8uL probe 15bT5 | 50,400 | 4,854 | 0.35, 0.2 | 126/160 | detection due to lower Ratio R=3 | | | | | small RNA H1 miR-15b > 16,800 | |
| nonew | | 2,863 | 0.35,005 | 92/112 | | | | | | **small RNA H1 miR-15b =** | **16,100** |
| | | | | | | | | | | | 0.06 |

#### Table S2, continues

| Flow cell, sample | probe copies | total events | (Ir/Io)max | working channels /active pores | Result/Conclusion | miRNA copies | Fig # AVE/RSD |
|---|---|---|---|---|---|---|---|
| **FAS88280** | | | | | | | |
| 3uL small RNA H2 | | 83,395 | 0.35<0.4 | 502/1516 | | | |
| nonew | | 75,846 | 0.35<0.4 | 479/1329 | | small RNA H2 miR-15b < 14,000 | |
| 3uL smRNA H2 + 4uL probe 15bT5 17.5fM | 42,000 | 138,200 | 0.2=0.3>0.35=0.4 | 445/1136 | detection | | |
| nonew | | 43,046 | 0.4,0.35 | 403/938 | | small RNA H2 miR-15b > 12,600 | |
| 3uL smRNA H2 + 3.6uL probe 15bT5 | 37,800 | 28,649 | 0.4>0.35 | 346/714 | silencing | **small RNA H1 miR-15b =** | **13,300** |
| nonew | | 20,325 | 0.4>0.35>0.2 | 235/422 | | | 0.07 |
| **FAS89359** | | | | | | | |
| buffer test | | 114,502 | 0.4+0.1,0.2,0.25 | 500/1526 | | | |
| nonew | | 126,724 | 0.2,0.4 | 496/1473 | | | |
| 3uL small RNA H1 | | 84,853 | 0.4 | 491/1429 | | | |
| nonew | | 142,416 | 0.2,0.4 | 464/1207 | | | |
| 3uL small RNA H1 | | 77,992 | 0.4>0.25,0.35 | 435/1079 | | | |
| nonew | | 94,426 | 0.4>0.45>0.2 | 362/841 | | | |
| 17.5fM | 50,400 | 82,990 | 0.25 | 321/682 | detection due to material shift to early Ir/Io | **small RNA H1 miR-15b < 16,800** | |
| nonew | | 141,939 | 0.25 | 246/484 | | | |
| **FAS90502** | | | | | | | |
| buffertest | | 163,655 | 0.4 | 505/1612 | | | |
| nonew | | 151,481 | 0.4 | 504/1571 | | | |
| use 4ul totRNA H4 | | 117,223 | 0.4>0.35 | 498/1536 | | | |
| nonew | | 119,697 | 0.4>0.35 | 473/1314 | | | |
| 4ul totRNA H4 + 4uL 17.5fM probe15bT5 | 42,000 | 133,952 | 0.2>0.35,0.4 | 460/1215 | detection, more events and low (Ir/Io)max | total RNA H4 miR-15b < 10,500 | |
| nonew | | 206,150 | 0.2, 0.35 | 416/999 | | | |
| 4ul totRNA H4 + 3uL 17.5fM probe15bT5 | 31,500 | 196,817 | 0.2, 0.35 | 379/862 | silencing due to less events at Ir/Io=0.35 | total RNA H4 miR-15b > 7,875 | |
| nonew | | 176,788 | 0.2, 0.35 | 320/654 | for both runs | **total RNA H4 miR-15b =** | **9,188** |
| | | | | | | | 0.20 |
| **FAS98320** | | | | | | | |
| buffer test | | 79,149 | 0.4 | 510/1521 | | | |
| nonew | | 88,827 | 0.4 | 501/1470 | | | |
| 1.5uL small RNA H2 | | 77,726 | 0.4 | 493/1394 | | | |
| nonew | | 97,350 | 0.2, 0.4 | 456/1098 | | | |
| 2uL totalRNA C1 | | 63,291 | 0.4,0.3,0.25 | 417/912 | | | |
| nonew | | 95,727 | 0.4>0.25,... | ND | control | total RNA C1 miR-15b > 21,000 | |
| 2uL total RNA C1 +4uL probe15bT5 17.5fM | 42,000 | 48,990 | 0.4>0.2,0.25 | 245/398 | silencing, half of the events | | |
| nonew | | 19,141 | 0.35 | 129/180 | | total RNA C1 miR-15b < 22,703 | |
| 1.85uL total RNA C1 +4uL probe15bT5 | 42,000 | 12,279 | 0.35 | 106/125 | detection, material shifts to early Ir/Io | **total RNA C1 miR-15b =** | **21,852** |
| nonew | | 5,308 | 0.25 | 78/85 | detection confirmed | | 0.06 |
| **FAS89319** | | | | | | | |
| buffer test | | 133,293 | 0.35>0.4 | 508/1652 | | | |
| nonew | | 127,415 | 0.35>0.4 | 508/1623 | | | |
| 4ul totRNA BRE CAN9 | | 222,511 | 0.35>>0.15 | 501/1536 | | | |
| nonew | | 92,794 | 0.35 | 486/1366 | | | |
| 4ul totRNA BRE CAN9 + 4uL probe15bT5 | 42,000 | 111,805 | 0.35, 0.2 | 463/1163 | detection, more events and material shift to Ir/Io=0.2 | total RNA Breast CAN9 miR-15b < 10,500 | |
| nonew | | 76,485 | 0.35 | 420/914 | | | |

For probe sequence, see Table 1.

PA, BRE and PRO stand for pancreatic, breast and prostate cancer sample, respectively; smRNA stands for small RNA and when the source is not identified it is small RNA from H6914 ($1^{st}$ lot) and when identified as (N/N) it is from H6914 ($2^{nd}$ lot). "Nonew" stands for a second 45min run with no added sample, and nonew2 for a third 45min run. The concentration of the stock solution as well as the number of OsBp moieties on the probe is identified in the first sample and applies to the rest of the samples with the same probe. Probe copies calculated from (probe concentration in fM) x (probe aliquot in $\mu$L) x 600. Total events as obtained using the *OsBp_detect* parameters listed in Methods under Workflow/Protocol. $(Ir/Io)_{max}$ are the maxima observed with the histogram of the total events (see Figures). Working channels/active pores, are reported by MinKNOW when the experiment is initiated. There are 2018 total nanopores, but only 512 channels, and therefore the number of working channels is reduced slower during the initial compared to the later experiments. Instead of determining a miRNA copy number, the result here is a YES/NO answer, i.e., whether the target miRNA is more or less than a certain copy number. To show how close to a miRNA copy number one can get, we determined an average value from the two experiments, one of detection and one of silencing, that lie closed to each other, and determined RSD which are between 7% to 16%.
